# SPROUT: A User-friendly, Scalable Toolkit for Multi-class Segmentation of Volumetric Images

**DOI:** 10.1101/2024.11.22.624847

**Authors:** Yichen He, Marco Camaiti, Lucy E. Roberts, James M. Mulqueeney, Eleftherios Ioannou, Marius Didziokas, Anjali Goswami

## Abstract

The segmentation of fine-grained and complex structures from volumetric data, such as 3D biomedical images, is a manually intensive process, with performance hindered by limited training data and the difficulty of adapting AI models for specialised datasets. Here, we introduce SPROUT, a user-friendly and interpretable segmentation framework that leverages domain-specific priors, enabling experts to translate their knowledge into reproducible, high-quality segmentations across diverse imaging modalities without the need for training data. Its adaptive design facilitates parameter transferability and improves generalisation across similar datasets. Implemented as scripts and a napari plugin, it supports interactive editing and scalable batch processing, lowering the technical barrier for domain experts.

We applied SPROUT to datasets spanning different imaging modalities, anatomical complexities, postures, and target structures, producing high-quality segmentations across 2D and 3D tasks. Quantitative comparisons with other methods on a representative dataset showed SPROUT achieved results comparable to expert-corrected interpolation while requiring substantially less manual input. In scenarios where SPROUT achieved high-quality results, supervised models often struggled to reach similar accuracy, highlighting the challenge of deep learning methods in complex domains. We also explored integration with foundation models to accelerate segmentation in high-contrast datasets, illustrating potential for hybrid workflows.

## Introduction

Three-dimensional (3D) or volumetric greyscale image data, such as those obtained through computed tomography (CT), micro-CT, and magnetic resonance imaging (MRI), are increasingly critical components of biomedical and biological research. These imaging modalities enable the non-destructive visualisation of complex internal structures in three dimensions, facilitating investigations of normal and abnormal morphology, anatomy, and tissue organisation (Hussain et al., 2022; Metscher, 2009). In biological systems, such scans often contain multiple regions of interest (ROIs), each corresponding to anatomically or functionally distinct components, which are essential for tasks such as 3D phenotyping, comparative anatomy, and developmental studies (Goswami et al., 2022). In medical contexts, volumetric images are widely used for diagnostic imaging, surgical planning, and treatment monitoring (Filippi et al., 2016; Thiesse et al., 2010). Beyond life sciences, volumetric imaging is also applied to other domains such as material science and industrial quality control (du Plessis & Boshoff, 2019; Moini et al., 2018).

Rapid acceleration in the digitisation of volumetric images, including those from biological and medical domains, is making large-scale, high-resolution datasets increasingly accessible to researchers (Altenkirch et al., 2008; Hashem et al., 2018). Available data have also expanded through open-access 3D libraries of bioimages and natural history specimens (Boyer et al., 2016; Marcus et al., 2007). Given this increasing availability of large-scale 3D datasets, accurate and efficient segmentation is becoming ever more critical. Regardless of context, segmentation enables the isolation of fine-grained, multi-class, domain-specific regions of interest and is essential for downstream analyses such as trait extraction, mesh reconstruction, precision measurement, and quantitative comparison.

Manual segmentation is labour-intensive and error-prone. These issues have been effectively addressed through artificial intelligence (AI) and deep learning for many applications. A wide range of deep learning models has been developed for various image modalities. Examples include convolutional neural network (CNN)-based architectures such as Mask R-CNN (K. He et al., 2017) and Deeplab (Chen et al., 2018) for natural images and U-Net and its transformer-based or 3D variants for biomedical volumetric data (Çiçek et al., 2016; Hatamizadeh et al., 2022; Ronneberger et al., 2015). Despite their success, these methods require training or fine-tuning and may not perform well in domain-specific tasks, particularly when anatomical knowledge is involved, or when fine-grained segmentation of multiple adjacent structures within a scan is required (Panayides et al., 2020; Tajbakhsh et al., 2016). It can be more challenging when dealing with various specimen shapes or postures, weak boundaries, or low contrast (Y. He et al., 2024). In such cases, creating a robust training set can be difficult, and the results may not meet expert expectations even after training. Furthermore, while various methods and techniques have been developed to enhance performance, the inherent lack of interpretability and the ‘black box’ nature of these models often make it difficult to further effectively fine-tune or optimise models in challenging domains (Hassija et al., 2024; Salahuddin et al., 2022).

Recently, foundation models have emerged as a new paradigm for segmentation tasks. These are large-scale models trained on diverse datasets to learn general-purpose representations and can be adapted to specific tasks through flexible prompts. Models such as the Segment Anything Model (SAM) are pre-trained on vast collections of real-world RGB images and support prompt-based segmentation (e.g., using points or boxes). These models generalise well to new datasets with little or no additional training, particularly for real-world RGB images (Kirillov et al., 2023; Ravi et al., 2024). To overcome limitations in biomedical imaging, domain-specific foundation models such as MedSAM and MicroSAM have been introduced (Archit et al., 2025; Ma et al., 2024). These models are trained on large greyscale datasets from medical and microscopy contexts and show improved performance in tasks such as organ and cell segmentation. Users can interactively refine segmentations by adding or adjusting prompts, making them highly adaptable in practice. However, most existing foundation models have been developed and evaluated within clinical or microscopy contexts. Their application to other settings—such as fine-grained segmentation tasks involving weak structural contrast, variable morphology, or changing posture in volumetric data—remains underexplored. These challenges are particularly common in non-clinical biological imaging, such as of natural history specimens.

Whether one adopts a traditional deep learning pipeline or uses foundation models, the initial step still involves manual annotation or pre-processing. Alternative approaches based on classic computer vision techniques can offer semi-automated workflows that reduce manual effort while maintaining accuracy, particularly when training data is unavailable or limited. These approaches often leverage priors, such as spatial priors (e.g., locations) or intensity priors (pixel values), and combine them with thresholding, interpolation, region growing, watershed, or other heuristic algorithms (e.g., blob detection) to generate accurate segmentations with reduced manual effort (Adams & Bischof, 1994; Bradski, 2000; Kong et al., 2013; Najman & Schmitt, 1994; Otsu, 1979). Several tools have successfully streamlined such methods into practical solutions. For instance, Biomedisa uses intelligent interpolation via weighted random walks to propagate user-defined segmentations across slices (Lösel et al., 2020), ilastik integrates interactive watershed and pixel classification for segmentations (Berg et al., 2019), and BounTI applies heuristic thresholding and region growth for 3D segmentation tasks (Didziokas et al., 2024). In addition, proprietary software such as AVIZO (Thermo Fisher Scientific) and Dragonfly (Object Research Systems) provide versatile toolkits for segmentation using both classic computer vision and deep learning techniques.

To address limitations in segmenting fine-grained, multi-class structures from volumetric data—particularly when expert-annotated training datasets are unavailable, boundaries are weak, or morphology is highly variable—we present **S**emi-automated **P**arcellation of **R**egion **O**utputs **U**sing **T**hresholding (SPROUT). SPROUT is a training-free and interpretable framework that leverages domain-specific priors and adaptive parameter search to separate and recover target ROIs in a controlled, consistent manner. It supports both fully batch processing and semi-automated workflows, with a graphical user interface (GUI) that enables specialists to produce reliable segmentations without advanced computational skills. In the sections that follow, we apply SPROUT to diverse imaging modalities and domains, compare it against alternative approaches, and explore its integration with segmentation foundation models for hybrid workflows.

## Results

### Overview of the SPROUT segmentation workflow

SPROUT is a segmentation framework originally developed for micro-CT scans of bones from natural history specimens and later generalised to a broad range of volumetric imaging tasks. It uses structural organisation and greyscale intensity patterns to perform segmentation without training data. This makes it effective for segmenting morphologically or texturally distinguishable structures, even under low-contrast conditions, and particularly suited to tasks involving morphologically complex objects where multiple parts need to be separated but large annotated datasets are unavailable.

The workflow consists of two main steps (Fig. 1a).

**Figure 1.**
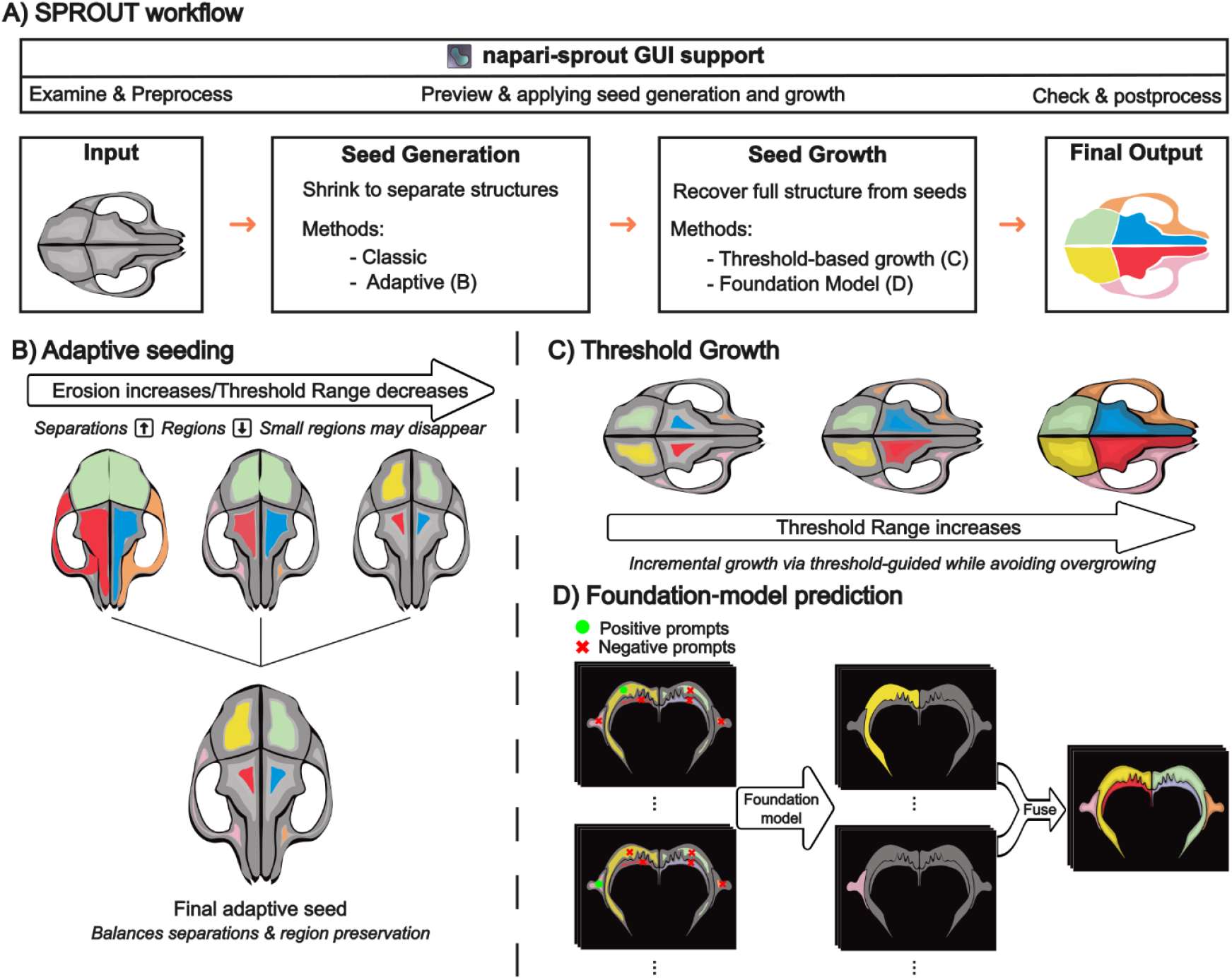
Overview of the SPROUT segmentation workflow. (a) The main pipeline comprises seed generation and growth, with GUI support via napari. (b) Adaptive seed generation compares seeds from multiple configurations to identify ROI splits and losses, producing a robust final seed. (c) Threshold-guided growth incrementally expands regions while avoiding overgrowth. (d) An alternative segmentation method uses positive (green dots) and negative (red crosses) prompts, derived from seed masks for each class in each slice, for the foundation model to make slice-wise predictions, which are then fused into a 3D output.

i. In seed generation, the image is decomposed into fine-grained candidate regions by shrinking structures to increase separation between adjacent components. To avoid over-reliance on a single parameter set, SPROUT includes an adaptive seed generation strategy that explores multiple threshold and erosion configurations, merging outputs to balance structure separation and preservation (Fig. 1b).
ii. In seed growth, the initial seeds are expanded to recover the full extent of each structure, either using an incremental threshold-guided method (Fig. 1c) or via SproutSAM (Fig. 1d), which applies segmentation foundation models for prompt-based volumetric prediction.

SPROUT’s design allows users to adjust separation and growth parameters that are directly linked to image features and target ROIs, making the process interpretable and reproducible across diverse modalities. While all functions can be executed via scripts, SPROUT is also available as napari-sprout, a plugin for the open-source 3D image viewer and analysis platform napari (Sofroniew et al., 2022). The plugin offers interactive parameter configuration, real-time visualisation of intermediate results, and region-level editing. Detailed implementation is described in the Methods section.

### Batch Segmentation with Adaptive Seeds

In datasets where objects share similar structural characteristics and are imaged under comparable conditions, separations between target regions can often be achieved across a range of threshold or erosion parameters, rather than relying on fixed values. SPROUT’s adaptive seed generation takes advantage of this property by exploring multiple parameter configurations and combining their results to generate robust, high-quality seeds. These final seeds can then proceed directly to the growth step. Notably, the seed growth process itself can also be guided by a range of thresholds, allowing incremental and controlled expansion. Together, these features make SPROUT particularly well suited for scalable and batch-friendly workflows.

#### Example 1: CT scans of Domestic Dogs (varied postures and cropping)

We demonstrate this approach on a set of CT scans of domestic dogs (*Canis familiaris*) produced at the University of Liverpool Small Animal Teaching Hospital. This is part of an ongoing study investigating the shape and morphological integration of limb bones in canids, requiring individual segmentation of these elements. This dataset has multiple characteristics that pose a challenge for AI-based segmentation methods: the scans have different voxel sizes and include different portions of individuals of different breeds in variable postures (Fig. 2). The greyscale values of corresponding anatomical regions remain consistent across the sample, with effective threshold values falling within a range. This enables the use of threshold-based adaptive seed generation across the dataset without requiring scan-specific tuning (for details, see Methods: Adaptive seed generation).

**Figure 2.**
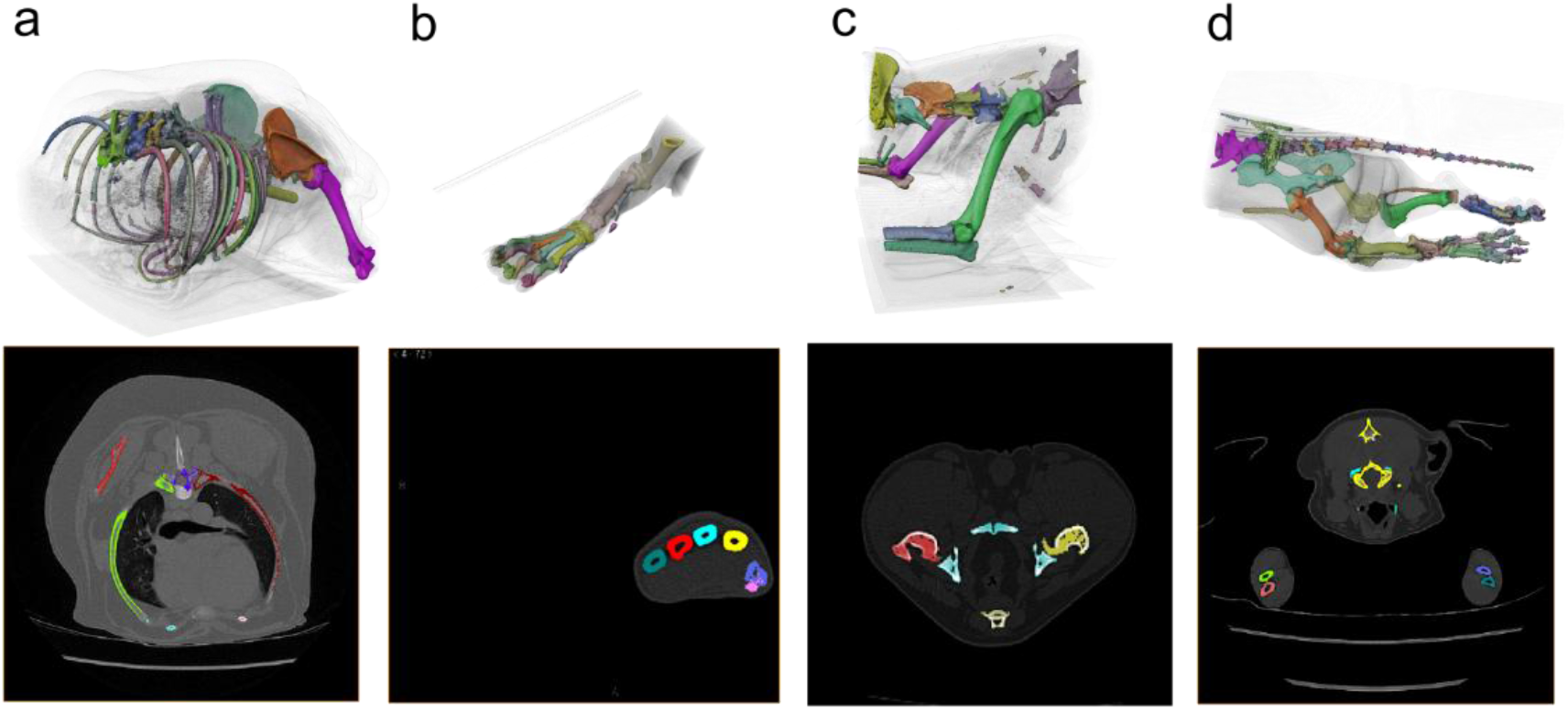
SPROUT segmentations of anatomical structures from CT scans of a domestic dog. (a– d) Four examples showing 3D visualisations (top row) and corresponding 2D slice segmentations (bottom row) for (a) thorax and shoulder, (b) lower forelimb, (c) shoulder, and (d) hindquarters including the tail. SPROUT robustly segments both large, complex regions (e.g. thorax) and small, low-contrast structures (e.g. tail vertebrae), demonstrating generalisation across anatomy and postures.

Despite this variability, the segmentations shown here were generated through the same batch process using the adaptive seed, followed by checking, correction and class mapping. As demonstrated, SPROUT can robustly capture the relevant structures across a diverse range of scans.

#### Example 2: Binary Mask Partitioning (variable numbers of regions)

SPROUT can operate not only on raw greyscale images but also on binary segmentation masks. Here, we applied SPROUT to the task of separating individual chambers within planktonic foraminifera from binary masks, as a part of a study investigating chamber ontogeny. The initial binary masks, representing the full chambered structure, were generated using a 3D U-Net trained from scratch with Biomedisa (Lösel et al., 2020), taking micro-CT scans of planktonic foraminifera as input. This model achieved an average Dice score greater than 0.95 (Mulqueeney et al., 2024). Separating these binary masks into individual chambers presents several challenges: (i) chambers are often connected due to calcification or limited resolution at their boundaries; (ii) the number of chambers varies across specimens; (iii) Chamber size and morphology can differ substantially within a single specimen. These properties make it difficult for deep learning, which typically requires a fixed number of classes for training.

Instead, we employ the erosion-based adaptive seed strategy in SPROUT (see Methods: Adaptive seed generation for details), as multiple thresholds for binary masks cannot be used. This approach applies morphological erosion incrementally to introduce separations between connected regions, while preserving small chambers that might disappear. The ability to balance separation and preservation makes adaptive seeding particularly well suited to this task. SPROUT was used for batch processing multiple specimens. As shown in Fig. 3, the method reliably separates individual chambers, even when the number and size of chambers vary between specimens.

**Figure 3.**
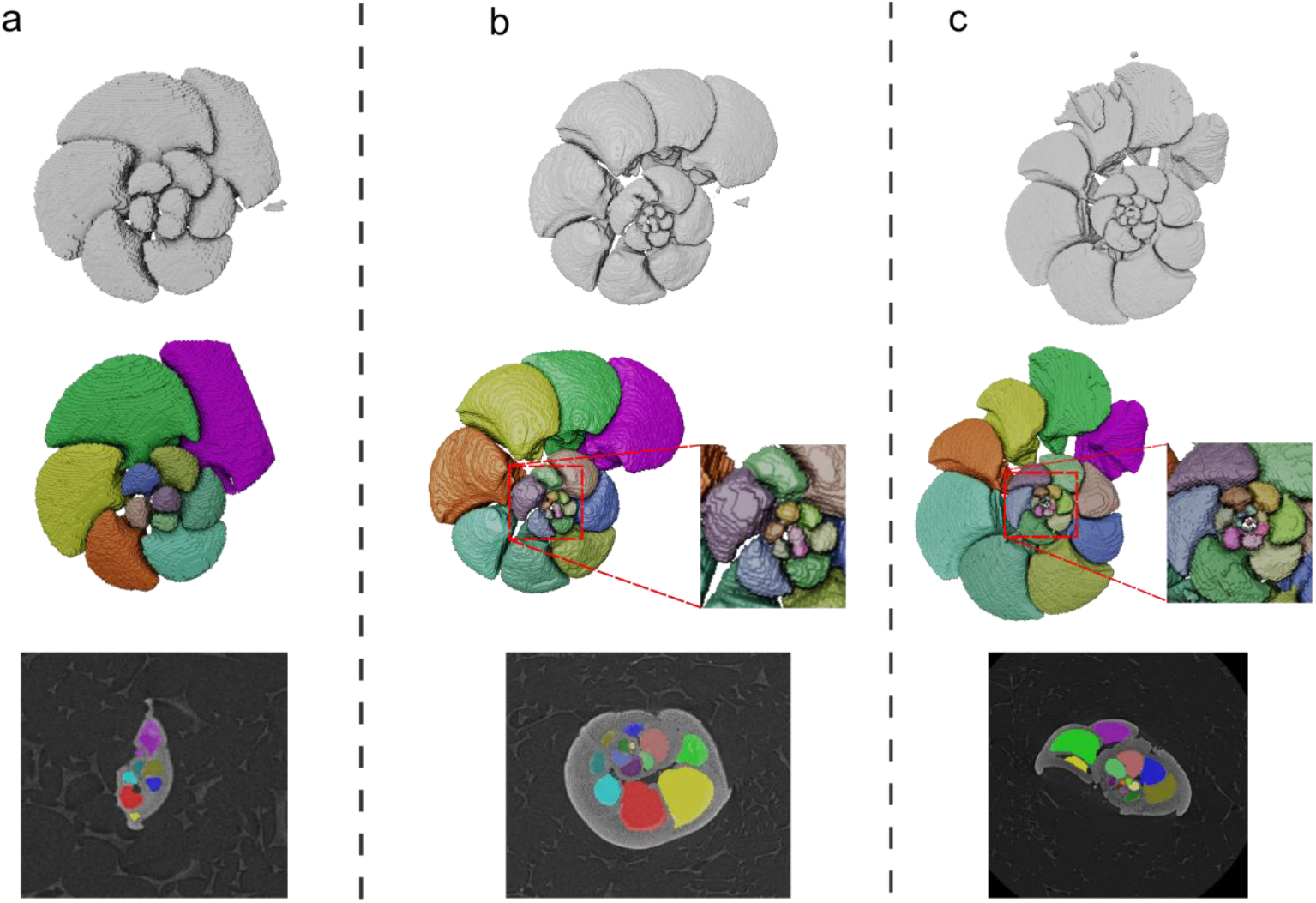
SPROUT segmentations of individual chambers from binary masks of planktonic foraminifera. (a–c) Specimens with 11, 20, and 29 chambers. For each example, we show (from top to bottom): the binary mask predicted by a trained 3D U-Net, the multi-label segmentation produced by SPROUT and a representative 2D slice showing the corresponding segmentation.

### Semi-automated Segmentation

#### Application for Complex Anatomical Structures

While SPROUT supports automated segmentation, some datasets pose anatomical or structural challenges requiring semi-automated workflows. One example is the segmentation of skulls into distinct bones from micro-CT scans, for a study aiming to compare the shapes of these bones across mammals. This task is difficult due to a combination of low contrast at sutural boundaries, complex interlocking cranial topologies and the large variation in bone sizes within a single specimen (Fig. a).

First, SPROUT supports the use of boundary masks to guide segmentation. In our example, we applied a morphological *top-hat* transform (Soille & others, 1999) and manually tuned its parameters to produce an intentionally exaggerated boundary mask (Fig. 4b). This boundary mask was then used explicitly as an input to SPROUT: during seed generation, it assists in splitting connected components more effectively; during seed growth, it constrains seed expansion, preventing overgrowth. For fine-scale recovery of the original structure, the boundary constraint can be temporarily removed in the final growth step, allowing small details to be restored (Fig. 4c).

**Figure 4.**
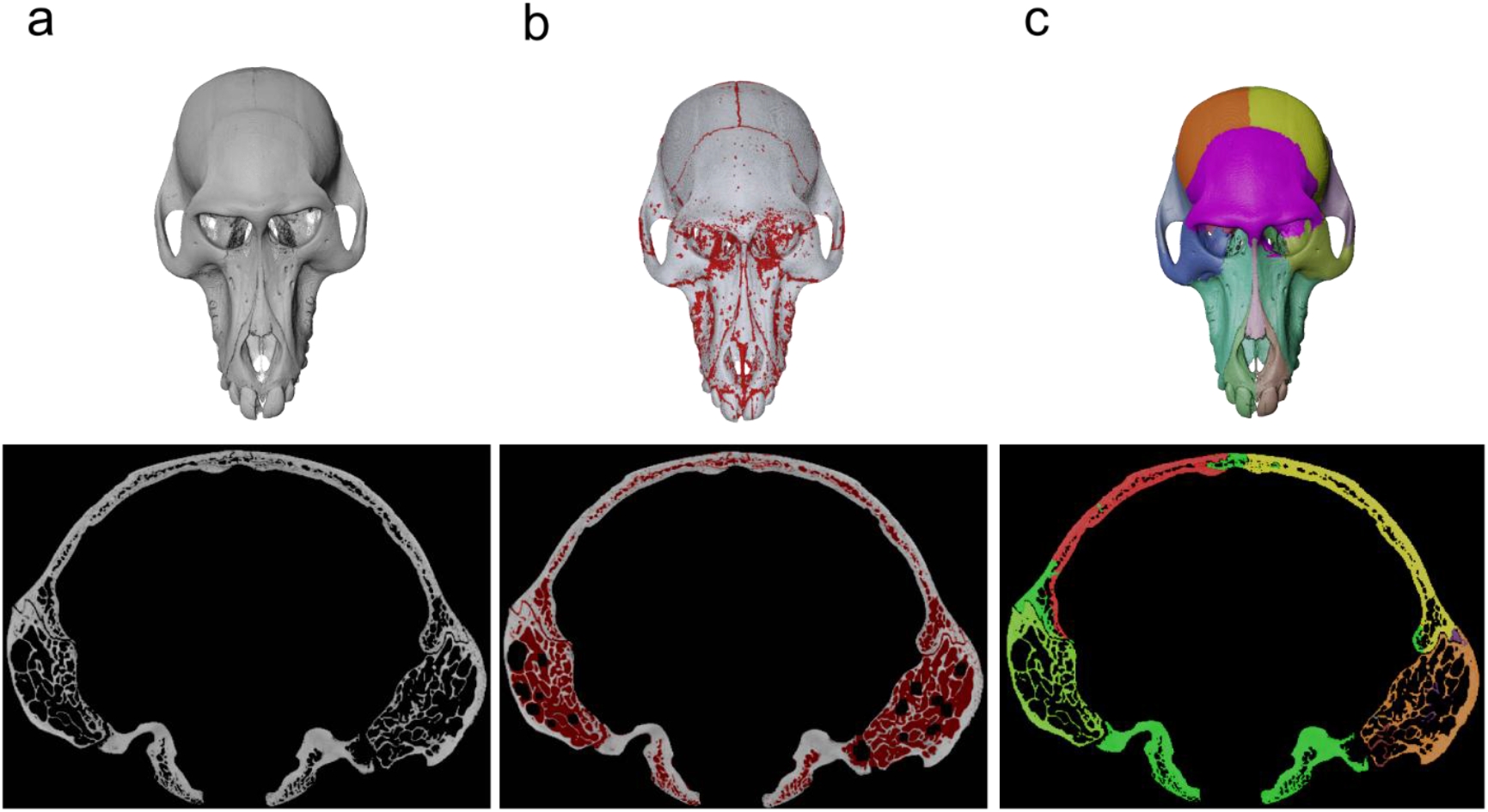
Semi-automated segmentation of a mammalian skull (Papio hamadryas) from micro-CT scans using SPROUT. (a) Original 3D rendering of the skull and a representative 2D slice from the micro-CT volume. (b) Boundary mask generated by applying a morphological top-hat transform with manually tuned parameters, intentionally exaggerating sutural boundaries to increase separations. The boundary mask is overlaid in red on both the 3D rendering and the corresponding 2D slice. (c) Final segmentation result after seed generation and growth in SPROUT in the 3D rendering and the corresponding 2D slice.

Second, manual refinement is supported via the napari-sprout GUI. Users can visualise seeds and apply region-level corrections, such as merging over-separated regions, removing irrelevant components, or splitting under-separated ones (see Supplementary Fig. 1c). These operations can be performed directly in 3D, making the correction process both intuitive and efficient. While pixel-level misclassifications, such as leakage into neighbouring classes, can occur, napari supports slice-by-slice pixel editing for these cases. In our experience, region-level correction is used far more often than slice-by-slice pixel editing for this task and provides more efficient refinement route.

#### Applications Across Modalities

We further assessed SPROUT on a diverse set of datasets spanning multiple imaging modalities and domains (Fig. 5). These comprised biological micro-CT and synchrotron CT datasets, including a skink (bones and dermal scales), mammalian skulls (individual cranial bones), and a Papuan weevil (tagmata and legs); microscopy datasets, including serial block-face electron microscopy (SBEM) of mouse retina (seven individual cells) and 2D confocal microscopy (all cells in the field of view); medical imaging datasets, including cardiac MRI (cardiac chambers and great vessels) and abdominal CT (vertebrae, ribs, kidneys, and the aorta); and an industrial CT dataset of concrete (aggregates and steel reinforcement). In all cases, SPROUT produced anatomically or materially coherent three-dimensional segmentations. Where necessary, results were refined using the napari-sprout interface, with the majority of corrections performed directly at the 3D regional level. For datasets outside our primary areas of expertise, segmentation outputs were optimised to the best of our ability but have not yet been formally validated by domain specialists.

**Figure 5.**
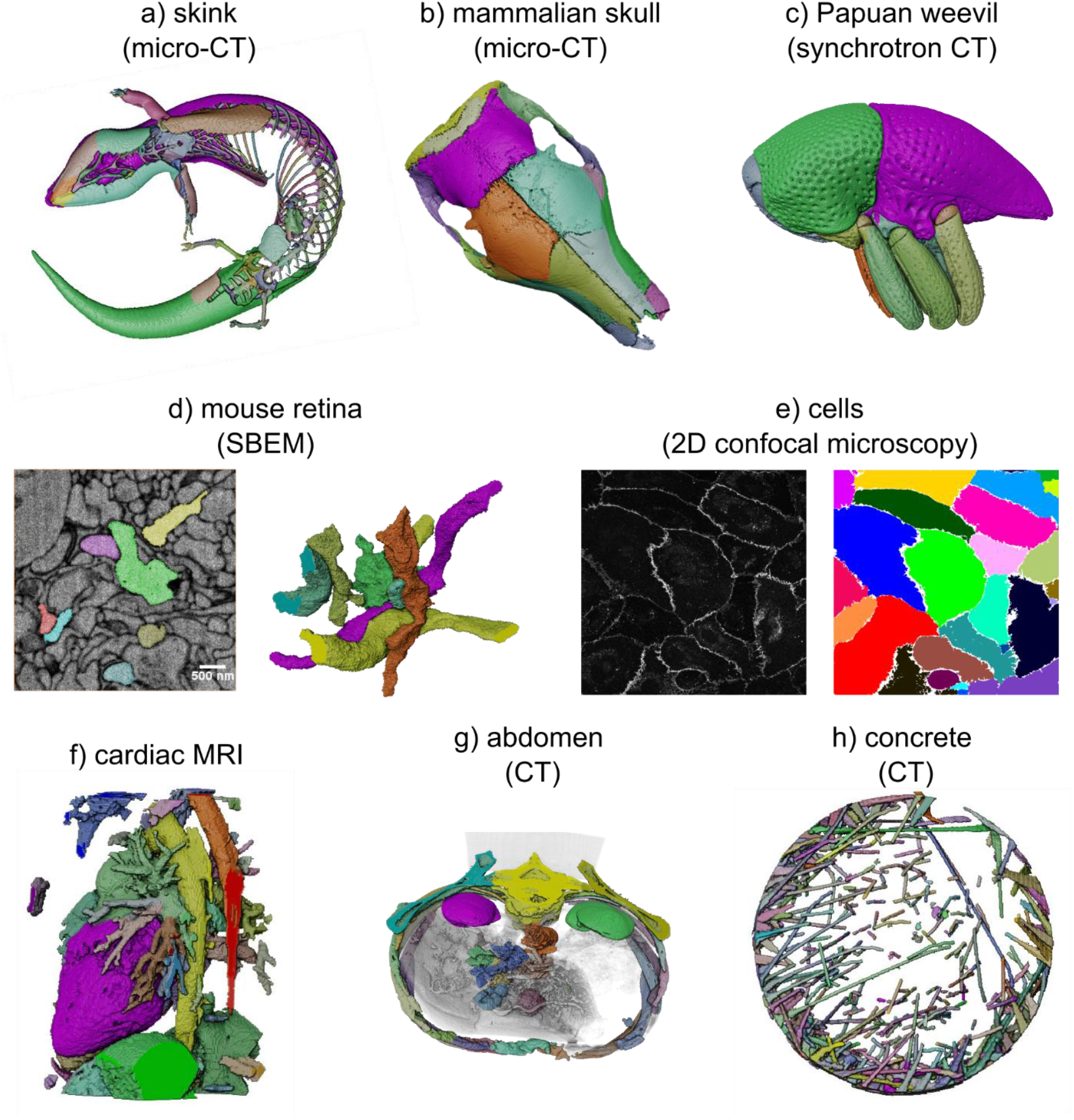
Applications of SPROUT across diverse imaging modalities and domains. (a) Micro-CT of a skink (Tiliqua scincoides), showing segmented bones and dermal scales. (b) Micro-CT of a mammalian skull (Orycteropus afer), segmented into individual cranial bones. (c) Synchrotron CT of a Papuan weevil (Trigonopterus sp.), segmented into tagmata and legs. (d) Serial block-face electron microscopy (SBEM) of mouse retina: 2D slice with overlaid segmentation (left) and 3D rendering of seven individual cells (right). (e) 2D confocal microscopy of cells: raw image (left) and segmentation (right). (f) Cardiac MRI, showing segmented cardiac chambers and great vessels. (g) Abdominal CT, segmented into vertebrae, ribs, kidneys, and the aorta. (h) CT of concrete, showing segmented aggregates and steel reinforcement.

### Comparison with Other Segmentation Methods

#### Interpolation and Deep Learning

To evaluate SPROUT’s performance against existing interpolation methods, we tested it on CT scans of nine specimens of the domestic dog (*Canis familiaris*), cropped to focus on the forelimb, and compared the results to those from Biomedisa’s smart interpolation. These scans varied in breed, voxel resolution, and limb posture. For consistent comparison, we selected four representative bones (scapula, humerus, radius, and ulna) across all specimens. Each specimen had every 10–50 slices annotated for interpolation. We used Biomedisa’s smart interpolation results, manually corrected slice-by-slice, as ground truth. This provided good accuracy with reasonable manual effort, while also avoiding evaluation biases that could arise from using SPROUT-generated labels.

We ran SPROUT using adaptive seed generation and automated batch processing across all nine scans. The resulting segmentations were filtered to retain only the four target bones for comparison. Pairwise Dice scores were calculated between (1) Biomedisa interpolation and ground truth, (2) SPROUT and ground truth, and (3) SPROUT and Biomedisa interpolation.

All three comparisons yield consistently high Dice scores, with average scores of 0.981 (GT vs. interpolation), 0.968 (GT vs. SPROUT), and 0.954 (interpolation vs. SPROUT). Although SPROUT’s overlap with the Biomedisa-based ground truth is slightly lower than that of Biomedisa interpolation, the differences are minor. Furthermore, in this task, SPROUT required less than 10 minutes of manual post-processing per scan. In contrast, preparing annotated slices for interpolation took ∼30 minutes of manual effort per scan and additional ∼5 minutes to check. Unlike interpolation, SPROUT also produced full multi-class segmentations, including non-target regions such as the ribs and mandible, without needing target-specific guidance (See Fig. 1).

We further evaluated deep learning performance on the same segmentation task using a 3D U-Net architecture implemented in Biomedisa. We used a separate dataset of 15 dog CT scans with SPROUT-generated and manually refined ground truth labels, with 10 scans for training and 5 for testing. Several model configurations were explored focusing on data augmentation, and they were trained until convergence, and summary results are shown in Table 1.

**Table 1.** Average Dice scores across test scans for 3D U-Net models trained with different configurations.

| Configurations | Dice Score |
| --- | --- |
| No augmentations | 0.641 |
| • Random flipping along X, Y, and Z axes.<br>• Random rotations 5 degrees<br>• Axis swapping | 0.712 |
| • Random flipping along X, Y, and Z axes.<br>• Random rotations 10 degrees<br>• Axis swapping | **0.745** |
| • Random flipping along X, Y, and Z axes.<br>• Random rotations 20 degrees<br>• Axis swapping | 0.742 |

Overall, we observed that data augmentation improved model accuracy, with the highest average Dice score (0.745) achieved using a combination of random flipping, 10° rotations, and axis swapping. In contrast, the model trained without augmentation reached only 0.641. We chose to use the Biomedisa 3D U-Net implementation because it is an accessible framework that has achieved good performance, achieving over 0.95 Dice score in the foraminifera chamber segmentation task described earlier. Detailed configurations and results are introduced in Supplementary Note 1.

We observed that augmentation improved overall performance, although gains varied across test cases. Lower-performing predictions often involved misclassification of pixels between anatomically distinct classes, rather than systematic under-segmentation or missing structures (Supplementary Fig. 2). These inconsistencies may reflect out-of-distribution challenges where certain test scans differ from training examples in anatomical shape, relative bone positioning, or image contrast.

Here, we chose this segmentation task for several reasons. First, the class count is fixed, making it easy to apply deep learning models. Second, the class count is small, which allows an acceptable time for manual process (e.g., labelling slices for interpolation). Finally, it is a realistic segmentation task: segmenting individual elements of a skeleton is a routine task in morphometric and biomechanical research in veterinary and medical science (Carman et al., 2022; Séguin et al., 2020).

#### ilastik Carving

We tested ilastik’s Carving module (Berg et al., 2019) and SPROUT on several datasets to evaluate their segmentation capabilities. Carving is a watershed-based interactive tool that requires users to manually annotate foreground and background regions on slices after computing a boundary map. A standard demonstration dataset provided by ilastik consists of a serial block-face electron microscopy (SBEM) volume of the mouse retina inner plexiform layer, to segment a single axon. SPROUT successfully segmented the target axon after using its intensity features. In contrast, ilastik’s result required refinement in several slices to prevent boundary leakage. Notably, while ilastik is designed to segment a single target object at a time, the same SPROUT run yielded multiple well-separated axons in addition to the target one. These results are illustrated and compared in Supplementary Fig. 3.

To further compare workflows, we applied the Carving tool to other scans used in this paper. When segmenting individual chambers from binary segmentations of planktonic foraminifera, annotating foreground and background for each chamber on a single slice was generally sufficient to produce reasonable results. However, achieving precise segmentations often requires additional manual refinement (see Supplementary Fig. 4). In contrast, when segmenting bones from CT scans of dogs, annotating a single slice typically yields only partial segmentation of the target bone. In addition, due to the unclear boundaries, segmentation can often leak into neighbouring regions, see Supplementary Fig.Although not quantitatively benchmarked, we tested multiple foreground/background annotation strategies, including varying slice positions, numbers and locations of annotations, and more aggressive versus more conservative markings, to ensure a fair evaluation of both workflows across diverse and representative imaging tasks.

### Evaluation of SproutSAM

We evaluated the segmentation performance of SproutSAM across three datasets (see Methods: SproutSAM for implementation details). The first comprised nine CT scans of dogs, with the task of segmenting four bone classes, as described in the interpolation and deep learning comparison section. The second consisted of ten binary segmentations of foraminifera chambers, where the goal was to identify individual chambers (with varying chamber numbers across scans). The third test is a 3D electron microscopy (EM) volume of the mouse retina (the same scan used in the ilastik evaluation section), with the objective of segmenting seven axon cells.

SAM is available in four architectural variants (tiny, base, large, and huge), each with increasing numbers of parameters. To assess performance across architectures, we tested five versions of SAM: the original ViT-B (base) and ViT-H (huge), and three models from MicroSAM (Archit et al., 2025) trained on domain-specific data: ViT-L-lm (large variant trained on light microscopy data), ViT-L-em (large variant trained on electron microscopy data), and ViT-B-medical (base variant trained on medical imaging). All three MicroSAM models used here represent the largest available variants for their respective domains. Each model was evaluated using seven different point-prompt sampling strategies. In the main text, we focus on model-level comparisons, with Table 2 reporting average Dice scores across all models and segmentation classes. Full results for all sampling strategies are provided in Supplementary Information (Supplementary Note 2 and Supplementary Fig. 6), which show that non-random strategies generally yield more accurate and robust segmentations across datasets and model weights.

**Table 2.**
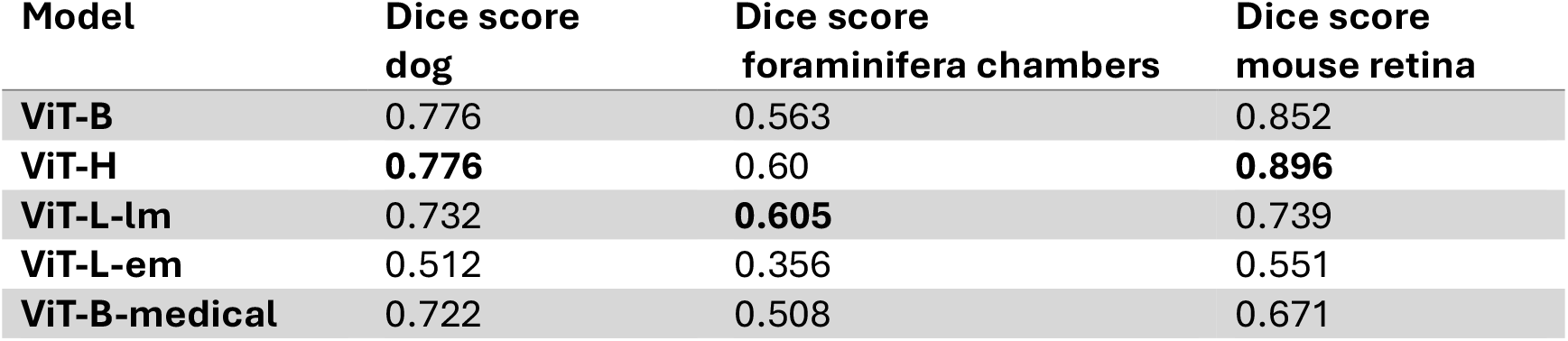
Average Dice score of foundation models across datasets.

| Model | Dice score<br>dog | Dice score<br>foraminifera chambers | Dice score<br>mouse retina |
| --- | --- | --- | --- |
| ViT-B | 0.776 | 0.563 | 0.852 |
| ViT-H | **0.776** | 0.60 | **0.896** |
| ViT-L-lm | 0.732 | **0.605** | 0.739 |
| ViT-L-em | 0.512 | 0.356 | 0.551 |
| ViT-B-medical | 0.722 | 0.508 | 0.671 |

Results indicate that the original SAM models performed best on the dog and EM datasets. Specifically, ViT-H achieved the highest Dice score on the EM volume (0.896), while both ViT-B and ViT-H achieved the top score on the dog dataset (0.776). For the foraminifera chambers, the ViT-L-lm model performed best (Dice score = 0.605), closely followed by ViT-H (Dice score = 0.600). The ViT-L-em model yielded the lowest Dice scores across all three datasets, including the EM task for which it was optimised (Dice score = 0.551). These results suggest that the pretraining domain of a foundation model does not necessarily guarantee optimal downstream performance. Overall, ViT-H demonstrated the most robust generalisation across tasks.

We further evaluated MicroSAM in its interactive mode using its napari plugin, which supports 3D propagation of segmentations from prompts placed on one or a limited number of slices. We qualitatively compared the performance of MicroSAM and SproutSAM across the test datasets, focusing on segmentation errors. Both tools can struggle on unclear boundaries, for example medullary cavities of the dog bones were incorrectly included.

SproutSAM tends to perform poorly when multiple target regions appear, leading to pixel misclassification or under-segmentation. These errors may stem from its strategy of sampling multiple classes per slice across three orthogonal axes, followed by fusion. However, this design can yield better 3D completeness. In contrast, MicroSAM’s interactive mode might need iterative prompt refinement, if the initial predictions were incomplete or failed to generalise across the volume. Users needed to add prompts on multiple slices. Its 3D propagation often failed when boundaries or spatial information were weak. Representative segmentation failure cases for both tools are shown in Supplementary Fig. 7. We did not include MedSAM in our evaluation. To the best of our knowledge, MedSAM is specifically trained for bounding-box prompts, whereas SproutSAM currently supports point-based prompts only.

Finally, we compared the runtime of SproutSAM with SPROUT’s original threshold-based growing approach. On larger scans, SproutSAM demonstrated improved speed. Detailed benchmarking results are provided in Supplementary Table 1.

### Computational Performance and Data Availability

We also evaluated the performance of SPROUT under multi-threaded execution, with results showing that parallel processing substantially reduces computation time (Supplementary Note 3, Supplementary Table 2, Supplementary Fig. 8). The datasets used for each evaluation and figure are summarised in Supplementary Table 3 and are available at Zenodo (https://zenodo.org/records/16857648).

## Discussion

Despite significant advances in segmentation tools, accurately and efficiently extracting individual structures from complex 3D images remains challenging. Fine-grained and multi-class segmentation tasks, scans with low contrast, irregular shapes, or variable topologies are difficult for both classical and deep learning methods, especially when annotated training data are time-consuming to produce or limited by dataset size. Under such constraints, fully supervised learning pipelines or general-purpose models may not yield satisfactory results, and heuristic methods may struggle to generalise across datasets.

SPROUT provides a lightweight and easily interpretable alternative for a wide range of segmentation tasks. It leverages greyscale intensities, morphological transformations (Soille & others, 1999), and structural priors in a modular workflow that does not rely on training data or specialised hardware, allowing users to tailor the segmentation process based on domain knowledge and image properties. SPROUT offers several core advantages that make it particularly suitable for these tasks.

First, it is designed with interpretability at its core. Parameters, including thresholds, erosion and dilation steps, and the number of components to extract, are explicitly defined and can be directly modified by users. This transparency stands in contrast to the opaque parameter spaces of many deep learning models and allows users to reason through segmentation logic based on the structural and intensity features of the image data.

Second, SPROUT supports efficient processing through a combination of adaptive seed generation and incremental seed growth. When applied to datasets with similar structural features, such as scans imaged under similar protocols, SPROUT can generate reliable segmentations using broad parameter ranges. This makes batch processing feasible without the need for tuning during the process. As with other segmentation methods, manual checking can be done at the final prediction stage.

Third, refinement in SPROUT is intuitive and efficient, often operating at the level of segmented regions rather than pixels or voxels. This enables users to perform most corrections directly in 3D, reducing the need for laborious slice-by-slice editing (e.g., fixing pixel misclassification or leakage). Intermediate results (e.g. partially grown segmentations) are interpretable and accessible and can serve as fallback outputs when they more closely match expert expectations. Users can readily inspect, merge, or split regions, as well as assign semantic classes, making the editing workflow more aligned with expert reasoning. These features are fully integrated as a GUI plugin into the napari platform. This lowers the technical barrier for users without programming experience and supports reproducible, visual, and interactive segmentation workflows.

Fourth, SPROUT is both adaptable and scalable across a wide range of image types and usage scenarios. As demonstrated in the results, it has been successfully applied to 3D volumetric data from CT, micro-CT, MRI, and electron microscopy, as well as to 2D microscopy images. Except for components designed for deep learning integration (i.e. SproutSAM), SPROUT relies solely on widely used Python libraries and runs efficiently on CPU, requiring only standard system memory. In contrast to deep learning methods, it does not require GPU acceleration, making it a cost-effective and accessible solution for researchers with limited computational resources. More importantly, as SPROUT’s core functions support both parallelisation and batch execution, the workflow can be efficiently scaled to large datasets and complex segmentation tasks, with substantial improvements in processing speed.

Building on these strengths, SPROUT has already been applied to collect the datasets presented in the Results section, including the segmentation of over 1,000 foraminifera specimens and more than 100 canine CT scans in batch, and over 100 mammalian skulls in semi-automated workflows. These fine-grained segmentations have substantially expanded available datasets in their respective domains, enabling new analytical opportunities. As these projects are still in progress, the datasets are not yet publicly available.

### Comparison with existing segmentation tools

SPROUT shares conceptual similarities with classical image segmentation methods such as blob detection and region growing, but differs in implementation details.

Blob detection methods, including those based on operators like Laplacian of Gaussian (LoG) and Difference of Gaussians (DoG), aim to identify blob-like regions in an image based on local intensity contrast and often apply connected component analysis to extract individual blobs. These approaches are typically used in microscopy and medical imaging to detect numerous small, round, or elliptical structures of similar size (e.g., cells or lesions) (Kong et al., 2013; Yu et al., 2014). While SPROUT’s seed generation also uses intensity and connected components to identify regions, it focuses on capturing structural or anatomical sub-regions within a larger object instead of blobs, using both intensity and morphological priors. Furthermore, compared to blob detection, SPROUT has adaptive seed strategies and parallelised processing pipelines implemented.

SPROUT’s growth step also resembles region growing algorithms (Adams & Bischof, 1994), as both expand from initial seed locations. However, classical region growing typically relies on user-defined seed points. In contrast, seeds from SPROUT’s seed generation step reflect structural separations, and its growth step supports multiple classes growing in parallel under incremental threshold-based guides. Compared to these methods, SPROUT provides a more integrated, modular, and scalable pipeline that generalises well to complex 3D segmentation problems.

In our evaluation, SPROUT achieved segmentation accuracy comparable to Biomedisa. Different from interpolation tools, which require manual annotations on representative slices for each scan, SPROUT is suitable for batch processing of structurally similar datasets by leveraging priors and adaptive seed generation. While this requires some understanding of the data’s structural and intensity characteristics, it can substantially reduce manual effort per scan, particularly in large-scale or multi-class applications.

Compared to supervised deep learning approaches, SPROUT requires no annotated training data and avoids the black-box nature of model inference, making it easier to inspect and refine segmentation outcomes. In our tests, deep learning models can produce pixel-level misclassifications, with different classes intermixed within the same slice (see Supplementary Fig. 2), and correcting such errors often requires slice-by-slice editing. In contrast, SPROUT’s threshold-based growth tended to produce more spatially coherent regions, with leakage occurring less frequently in our experience. Many corrections could therefore be performed as 3D region-level edits, which are generally more efficient. We tested multiple hyperparameter configurations to improve the deep learning results and observed gains from data augmentation, but performance eventually reached an optimisation plateau. Admittedly, further improvements could be achieved by tuning hyperparameters, testing alternative architectures, or adding more training data, but these require time and domain expertise in deep learning. Nevertheless, when optimised, deep learning models can often achieve high accuracy and offer a fully automated workflow.

We also compared SPROUT with ilastik’s Carving module, a watershed-based tool. Across the examples we tested, SPROUT consistently yielded more accurate segmentations, particularly in cases with ambiguous or low-contrast boundaries.

Finally, a feature of SPROUT is that it can produce a full, multi-class segmentation of the image based on global intensity and morphological information. This enables downstream selection or filtering of target regions post hoc. In contrast, many existing tools, such as supervised deep learning and interpolation, require users to predefine specific targets, e.g., through training datasets and selected slices.

### Integration with segmentation foundation models

Foundation models have recently emerged as a powerful paradigm for AI-based segmentation. While developing a volumetric foundation model is beyond this study’s scope, applying existing models to volumetric data can require the manual placement of prompts across many slices, which is laborious and may still fail to capture an object’s complete 3D topology. To address this, we developed SproutSAM as a framework for automated 3D prompt engineering, using SPROUT’s globally-aware, multi-class seed masks to construct point-based prompts across all relevant slices, replacing manual annotation.

In evaluation, SproutSAM performed particularly well on datasets with clear boundaries, such as the EM retina volume, where high-quality SPROUT seeds produced accurate predictions, often faster than SPROUT’s standard threshold-based growth. In more complex cases, such as micro-CT scans with segmentation leakage into adjacent structures, performance was limited. This reflects both a mismatch between our data and the available model weights and the inherent inconsistencies of reconstructing 3D objects from 2D prompts. Manual placement of prompts did not improve results in these cases either. SproutSAM thus represents a proof-of-concept for a hybrid workflow in which an interpretable method (SPROUT) supplies structural priors to guide a less-interpretable foundation model. This prompting approach could be extended to generate high-quality volumetric labels for fine-tuning existing models and training future robust, domain-specific 3D foundation models.

## Conclusion

SPROUT is more than an alternative segmentation tool; it establishes a new workflow for complex image analysis that combines interpretability, adaptability, and expert guidance in a single, modular environment. It requires no training data, and its inputs allow domain experts to directly leverage structural priors and greyscale intensity. By exploring parameter ranges and applying adaptive seed generation, users can efficiently batch process structurally similar datasets. SPROUT combines interpretability, efficient 3D region-level editing, and scalability, enabling researchers to generate accurate and reproducible segmentations across diverse tasks. In challenging tasks, when results from some interactive methods are not ideal, users often have limited control over how to refine them, due to low transparency or complex parameters. In contrast, SPROUT enables controlled re-runs on its outputs, so refinements remain interpretable.

Its integration as a napari plugin facilitates a seamless combination with other tools, such as MicroSAM, and supports workflows that incorporate foundation models. We have explored strategies for generating prompts from SPROUT outputs as inputs to these models. By uniting deterministic control, adaptive scalability, and expert-guided interactivity, SPROUT delivers a robust and extensible platform for large-scale, domain-specific image analysis, empowering researchers to achieve accurate, transparent, and reproducible results across diverse segmentation challenges.

## Methods

The SPROUT workflow and its main components have been summarised in the Results section (Fig. 1). Here, we provide a detailed description of each component, including seed generation, adaptive seed generation, seed growth, SproutSAM integration, and the napari-sprout GUI, along with implementation details.

### Seed Generation

The seed generation step aims to extract structurally meaningful and well-separated candidate regions from a greyscale image, based on a user-specified number of regions. This is achieved by first applying a thresholding operation to generate a binary mask, isolating the foreground (e.g., target structures) from the background. A morphological erosion can then be applied to the binary foreground, reducing its size and potentially increasing the separation between adjacent regions, particularly when boundaries are weak or narrow.

By adjusting parameters such as the threshold range (i.e., the lower and upper threshold bounds) and the number of erosion iterations, users can control the reduction of foreground regions in an interpretable manner. Following these steps, connected components analysis is used to separate the binary foreground into discrete objects, with an optional filtering step to retain the *n* largest components based on size.

### Adaptive Seed Generation

While varying threshold and erosion parameters can produce different candidate seeds, manually searching for the optimal configuration is time-consuming and may lead to suboptimal results. There is often a trade-off between achieving more separations and preserving smaller but meaningful regions. For example, a seed configuration that successfully splits a large structure may cause smaller elements to disappear due to excessive erosion.

To address this, we introduce an adaptive seed generation method that explores multiple configurations and merges information across them (Fig. 1b). Starting from an initial configuration, SPROUT iteratively generates a sequence of seeds where regions become smaller. At each iteration, it compares the new seed to the previous one, identifying: (i) Separations: regions that were previously connected, but now split into distinct components; and (ii) Disappearances: regions that existed before but have been eliminated. Each potential split can be validated using rules to determine whether it is a valid separation, helping to avoid over-fragmentation caused by noise or small artefacts.

To ensure that seed masks progressively decrease in size across iterations, SPROUT provides two adaptive strategies:

- **Erosion-based**, where the erosion is gradually increased under a fixed threshold.
- **Threshold-based**, where the threshold range is progressively narrowed under a fixed erosion.

The final adaptive seed typically outperforms any individual seed produced under a single configuration. It eliminates the need for precise tuning of input parameters by aggregating information across a range of erosion or threshold settings. Intermediate seeds generated during this process can also be saved and reused. While adaptive seed generation is more computationally intensive, it is particularly useful when no single parameter setting yields satisfactory separation.

### Seed Growing

The growth step aims to recover the full morphology of each target structure by expanding the seed regions produced during the previous step. Since these initial seeds are intentionally smaller than the actual components due to erosion or thresholding, they need to be grown back to their original size based on specified image thresholds.

SPROUT uses morphological dilation to expand each seed region iteratively. However, as dilation does not incorporate boundary information, it may lead to expansion beyond the target structures. To address this, the dilation process is often guided by threshold ranges, which restrict growth. Specifically, growth begins with a narrow range that allows for limited growth, and it is gradually widened until a final target threshold is reached (Fig. 1c). This strategy ensures that growth proceeds incrementally and stably, allowing structures to recover their full size while minimising the risk of overgrowth or leaking into adjacent elements or the background, particularly in cases where final thresholds alone do not offer clear separation between neighbouring regions. Like adaptive seed generation, intermediate results produced during this progressive growth process can also be saved.

### SproutSAM

In addition to Sprout’s threshold-based region growing approach described above, SproutSAM provides an alternative method for converting seed masks into final segmentations, leveraging segmentation foundation models such as SAM. These models generate 2D segmentation masks based on user-provided prompts, such as positive and negative points that indicate whether a location belongs to the target region.

To adapt these 2D models to volumetric data, SPROUT uses the 3D seed masks to generate slice-wise prompts along the X, Y, and Z axes. For each class, and on each slice along a given axis, we sample positive points from within the seed region and negative points from regions belonging to other classes. The model predicts segmentation masks per class and per slice. For example, given a 3D volume of size 100×50×20 with 3 segmentation classes, SproutSAM produces predictions for 300 slices along the X axis (100 slices × 3 classes), 150 along the Y axis, and 60 along the Z axis. These outputs are then aggregated across slices and axes to reconstruct the final volumetric segmentation. SproutSAM supports both SAM and SAM2 and allows custom-trained model weights. Details of prompt sampling strategies and fusion procedures are provided in Supplementary Note 4 and illustrated in Supplementary Fig. 9. Bounding box prompts are not supported in the current implementation, as SPROUT seed masks typically occupy only a subset of the full object extent. Bounding boxes derived from such seeds are often unreliable and may lead to inaccurate segmentations.

SproutSAM performs deep learning inference using PyTorch and is optimised for graphics processing unit (GPU) acceleration. Although it can operate on CPU, performance is improved on a GPU. GPU execution makes use of dedicated high-bandwidth video memory (VRAM), whereas CPU execution relies on standard system memory (RAM).

### napari-sprout

While all SPROUT functions can be executed via scripts and configuration files, we recognise that many users, particularly those with limited programming experience, benefit from being able to set parameters interactively and visualise intermediate results in real time. To support this, we developed napari-sprout, a graphical user interface (GUI) plugin integrated into the open-source napari viewer. The plugin allows users to load and save greyscale scans and segmentation masks, interactively adjust parameters, and directly apply core SPROUT functions, including seed generation, region growth, and post-processing. It also includes editing tools optimised for 3D use, enabling region-level operations such as merging or splitting components, class assignment, and quick inspection of results. Full GUI details and usage instructions are provided in Supplementary Note 5, and a visual demonstration is shown in Supplementary Figure 1.

### Implementation Details and Optimisation

SPROUT is implemented in Python (v3.10) and supports both script-based pipelines and GUIs. It is compatible with major operating systems (Windows, macOS, and Linux) and supports multi-threaded and batch processing for high-performance or cloud computing environments, making it suitable for large-scale dataset analysis. To facilitate integration with common analysis platforms, we also provide Avizo and ImageJ plugins. The implementation incorporates parallelisation and memory optimisation to ensure efficient handling of large 3D scans. Further details are provided in the Supplementary Information: Avizo and ImageJ/Fiji plugin integration (Supplementary Note 6), parallelisation and memory optimisation (Supplementary Note 7), and installation and usage instructions, including script-based examples (Supplementary Note 8 and Supplementary Note 9). For the latest updates, please visit the project repository (https://github.com/EchanHe/SPROUT). Unless otherwise stated (e.g., Biomedisa-based results), all evaluations in this study were performed on a workstation running Windows 11, equipped with an 11th Gen Intel(R) Core(TM) i7-11800H CPU @ 2.30 GHz, 48 GB RAM, and an NVIDIA RTX 3080 GPU (16 GB VRAM).

## Acknowledgements

We’d like to thank A. Sharp and E. Grisan for suggestions and technical insights, which helped the development of SPROUT. We would like to thank R. Fleuty, R. Foster and T. Weber from the Technology Solutions at Natural History Museum, London for providing help on computational resources. We thank T. Macrini and G. Pandelis for contributing the aardvark and the skink scans to Digimorph and Morphosource, respectively. Foraminiferal specimens were scanned at the µ-VIS X-ray Imaging Centre at the University of Southampton. Dog scans courtesy of Liverpool Veterinary School Clinical Collaboration Platform, with thanks to T. Maddox. We thank C. Cooney for supporting the development of this method. We also thank L.K. Phng and H. Wint for imaging and providing the confocal endothelial cell image.

## Funding information

This work was funded by Leverhulme Trust grant RPG-2021-424, NERC grants NE/S007210/1 and NE/P019269/1, Royal Society APEX grant AA21\100124, and BBSRC grant BB/X014819/1. Ethics approval for dog scans granted by University of Liverpool Veterinary Research Ethics Committee (VREC), approval codes VREC1311 (non-greyhound breeds) and VREC1339 (greyhounds).

## Author Contributions

YH conceptualised the project, developed the method, and wrote the manuscript. MC conceptualised the project and contributed to manuscript writing. MC, LR, and JM gathered data and tested the method. EI developed the method. AG conceptualised the project. All authors contributed to the development of the method and manuscript and reviewed and approved the final manuscript. All authors declare no conflict of interest.

## Notes

### Competing Interest Statement

The authors have declared no competing interest.

### Summary of Updates

Major Revisions in This Version New User Interface: Added a user-friendly graphical user interface (GUI) for SPROUT, implemented as a napari plugin. This greatly improves accessibility and interactivity, enabling a broader range of scientists to use the workflow effectively. Foundation Model Integration (SproutSAM): Introduced a new section exploring the integration of SPROUT with foundation models such as the Segment Anything Model (SAM). We present SproutSAM, a proof-of-concept workflow that uses SPROUT outputs to automatically generate prompts, bridging interpretable classical algorithms with state-of-the-art AI methods. Expanded Evaluation: Significantly extended validation, including more rigorous quantitative and qualitative comparisons against other leading segmentation methods across a wider range of volumetric datasets. Manuscript Revision: The manuscript has been substantially rewritten to incorporate these new features and results. Clarity and logical flow have been improved, new figures have been added, and much of the detailed technical content has been moved to Supplementary Information to make the main text more concise and accessible.

