## supplementary information for "SPROUT: A User-friendly, Scalable Toolkit for Multi-class Segmentation of Volumetric Images"

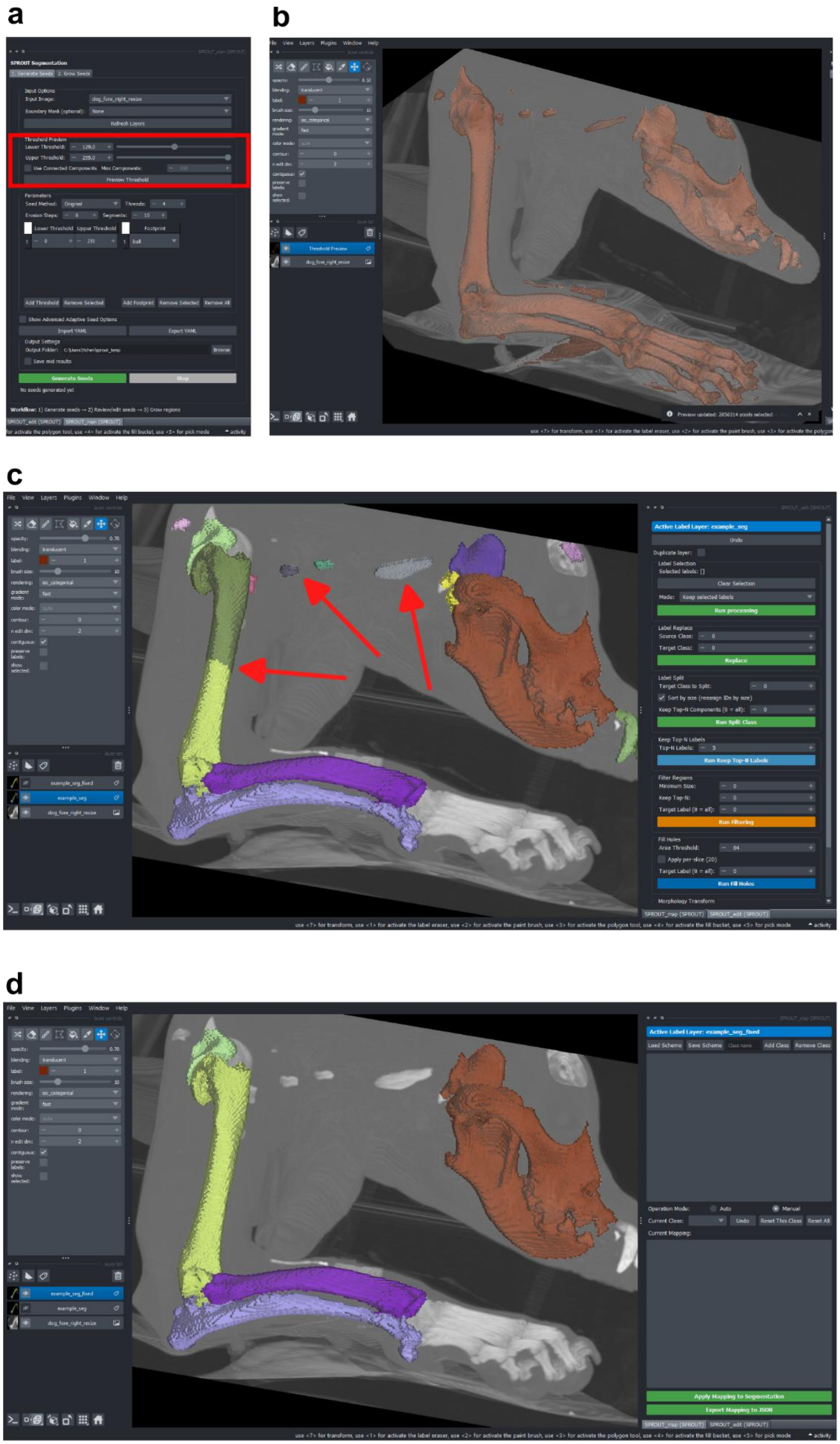
a
b
c
d

***Supplementary Figure 1. Key interface components of the napari-sprout plugin.*** *(a) Seed panel with parameter controls (e.g., threshold and erosion), and a preview sub-panel (highlighted in red) allows users to visualise thresholds before running the segmentation. (b) Threshold preview visualisation showing the segmented regions on a dog CT scan. (c) An example of segmentation result requiring post-processing, such as merging connected components and removing noise. The right panel shows the region-level editing interface for merging, removing, or reassigning segments. (d) Final refined segmentation (left) and the class mapping panel (right), which enables assignment of semantic labels to regions. Note: The grow panel, which shares a similar interface and parameter structure with the seed panel, is not shown here for simplicity. See Supplementary Note 5 for a detailed description of the napari-sprout implementation.*

### Supplementary Result

#### Supplementary Note 1: Comparison with deep learning

We used the Biomedisa platform (Lösel et al., 2020) to train a 3D U-Net (Ronneberger et al., 2015) for segmenting four bone classes from dog CT scans. Biomedisa employs a patch-based implementation of 3D U-Net with overlapping patches (patch size: 64x64x64 voxels, stride: 32). Besides configurations about augmentations we mentioned in the main text, training was performed using stochastic gradient descent with a learning rate of 0.01, decay of 1×10e-6, momentum 0.9. A batch size of 24 was used and 20% of training scans as the validation set, and models were trained for 50 epochs. We also tested alternative hyperparameter settings, but the above configuration yielded the most consistent and converged results.

While augmentation generally improved average performance (see main text), model accuracy varied substantially across individual test scans. For example, one scan consistently achieved high Dice scores across configurations, with the best model reaching 0.907. In contrast, another scan showed much lower scores overall, with the best result remaining below 0.517. These differences may reflect anatomical variation, imaging artefacts, or mismatches in posture or shape compared to training examples.

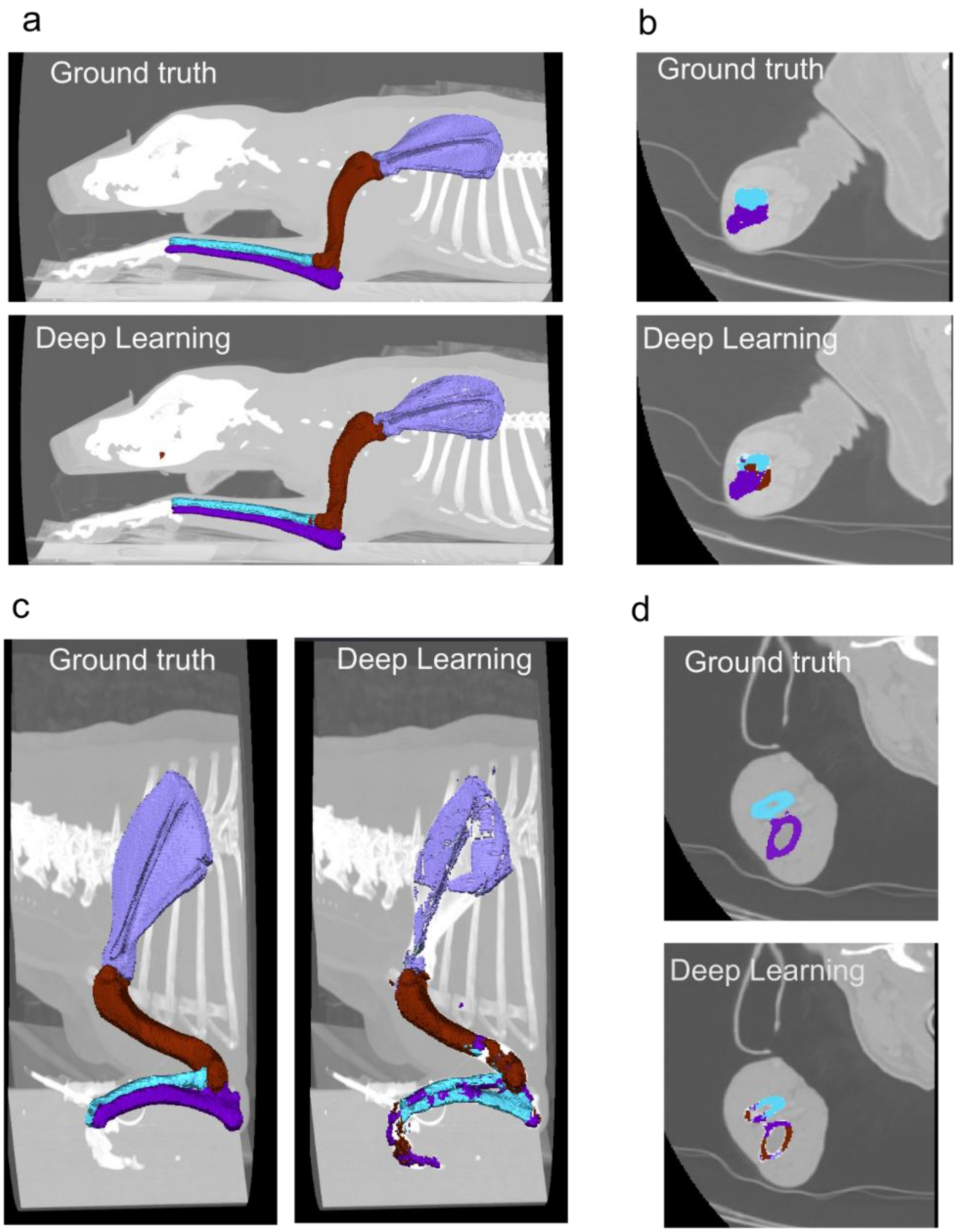


***Supplementary Figure 2 Comparison of deep learning segmentation results and ground truth segmentations for two representative cases.*** *(a, b) An example with a high Dice score (0.907). (a) The predicted structures closely match the ground truth. (b) A zoomed-in slice shows only minor misclassifications at structural junctions. (c, d) An example with a low Dice score (0.517). (c) Multiple structures are misclassified or merged. (d) A zoomed-in slice highlights the extensive pixel misclassification.*

#### Comparison with ilastik Carving

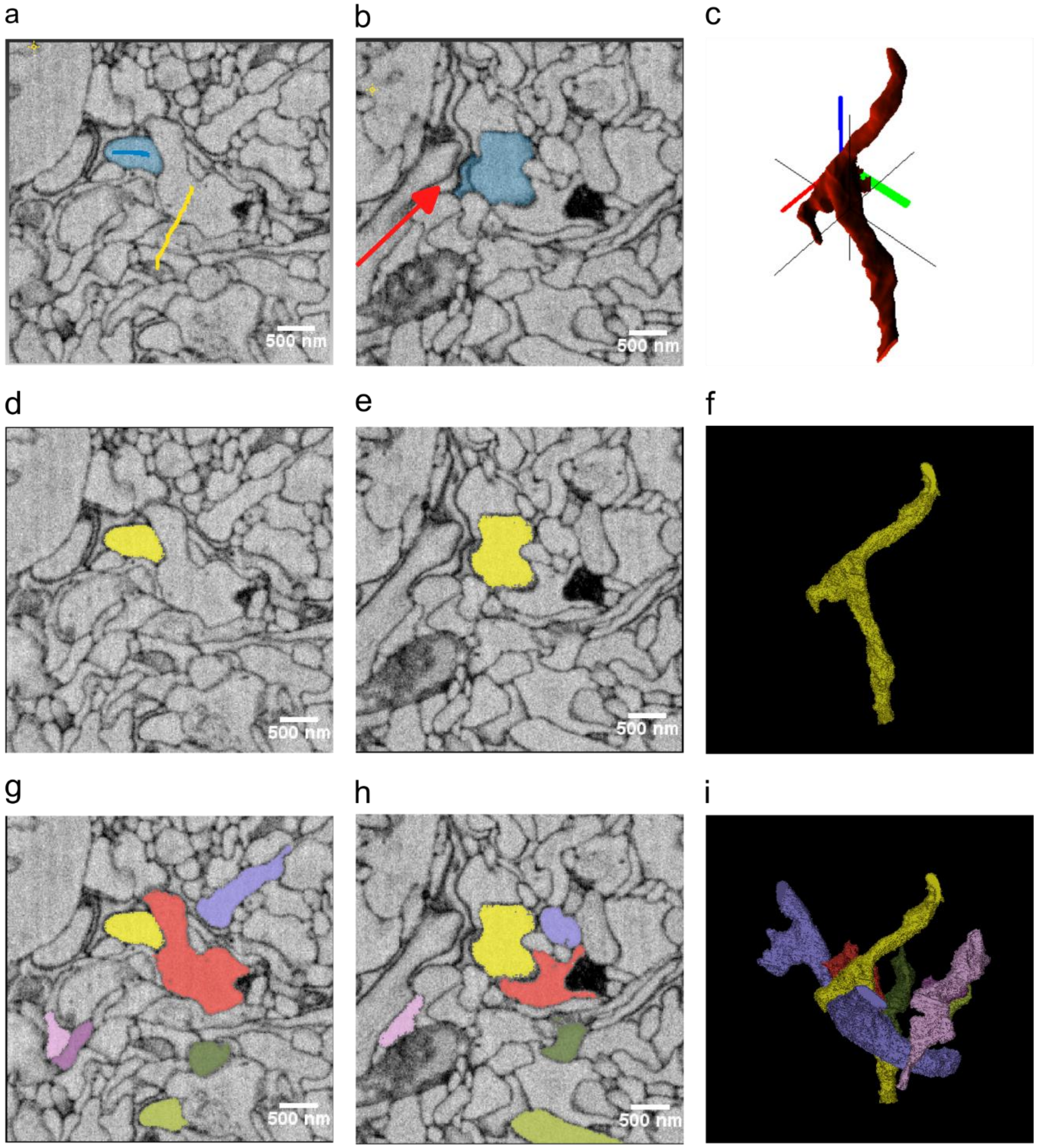


***Supplementary Figure 3. Comparison of SPROUT and ilastik Carving results using the example dataset from the official ilastik tutorial (https://www.ilastik.org/documentation/carving/carving).*** *The dataset is a serial block-face electron microscopy (SBEM) volume of the mouse retina inner plexiform layer, and the task is to segment a single axon. (a–c) ilastik results: (a) an example slice annotated with foreground (blue brush stroke) and background (yellow brush stroke) and the segmented region is in blue; (b) a slice showing segmentation leakage (indicated by a red arrow); and (c) the 3D rendering in ilastik. (d–f) SPROUT results, with the segmented region in yellow: (d) the*

*corresponding slice to (a), (e) the corresponding slice to (b), and (f) its 3D rendering. (g–i) Full SPROUT result showing multiple axons segmented in the same run, including corresponding slices and the 3D rendering.*

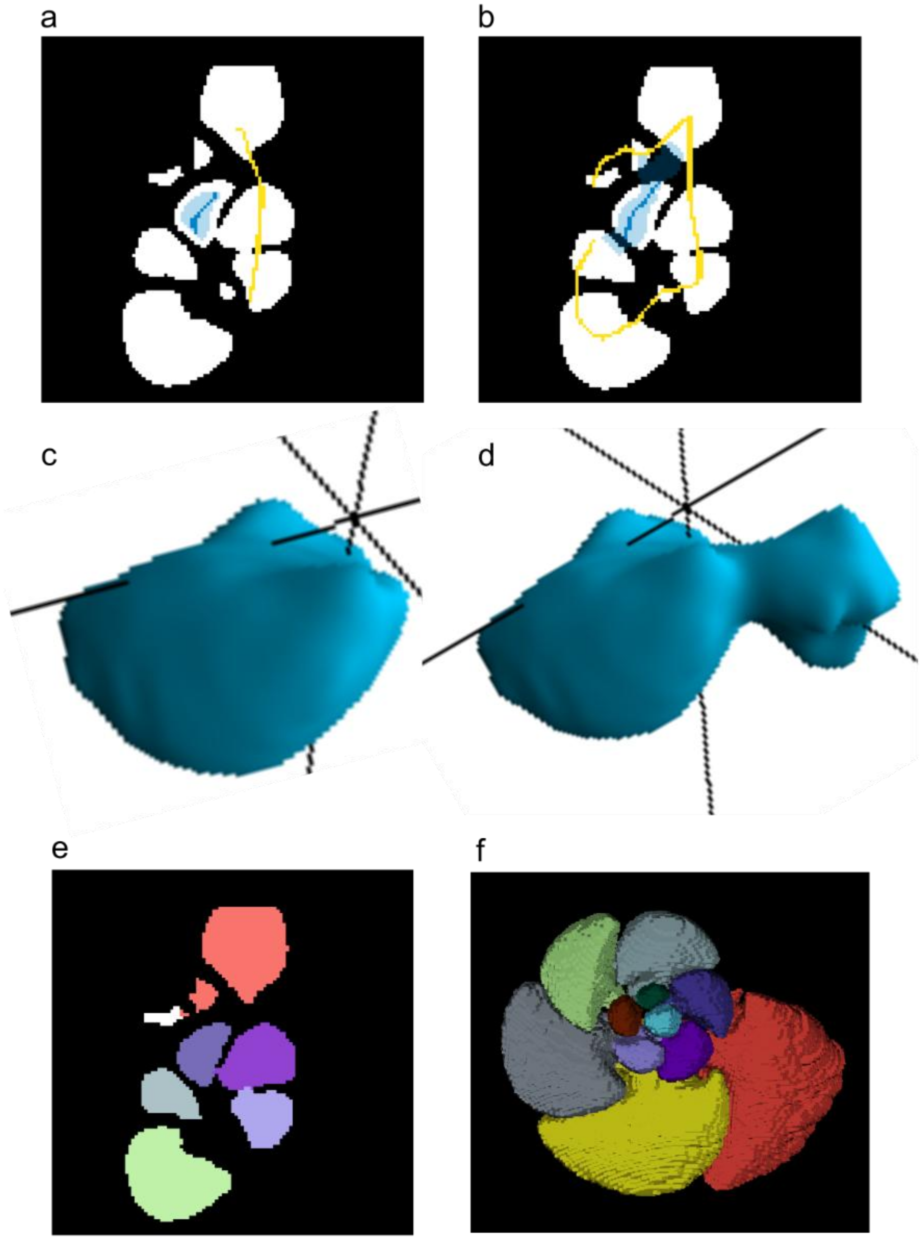


***Supplementary Figure 4. Comparison of ilastik carving and SPROUT for segmenting chambers of planktonic foraminifera.*** *(a–b) ilastik carving results on a single slice using foreground (blue brush strokes) /background (yellow brush strokes) configurations. In (a), the foreground is conservatively annotated, producing a chamber slightly smaller than its true boundary. In (b), a more aggressive annotation tightly covers the chamber and excludes other regions but causes leakage into neighbouring chambers. (c–d) 3D views of the ilastik segmentations showing the difference in volume and the leakage effect. (e) SPROUT*

*segmentation on the same slice and (f) the corresponding 3D result, generated without manual annotation, show accurate separation of the target chamber.*

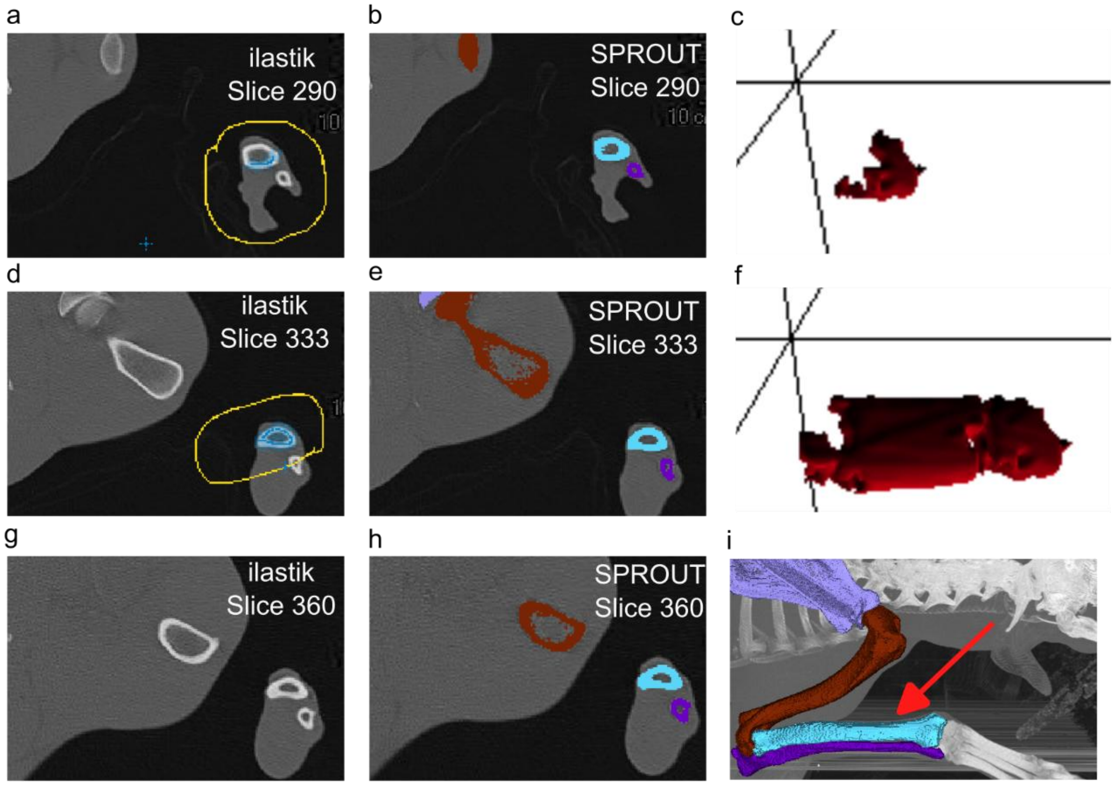


***Supplementary Figure 5. Comparison of ilastik carving and SPROUT for segmenting a bone in a dog CT scan.*** *(a) ilastik carving result with only slice 290 annotated. (b) SPROUT result on slice 290. (c) Corresponding ilastik 3D volume based on slice 290 annotation. (d) ilastik carving result after adding slice 333 as an additional annotation. (e) SPROUT result on slice 333. (f) ilastik 3D volume based on annotations from slices 290 and 333. (g) ilastik result on slice 360, showing no segmentation despite prior annotations. (h) SPROUT result on slice 360. (i) 3D rendering of the SPROUT result, with the target bone indicated by a red arrow. Foreground (blue) and background (yellow) annotations in ilastik were applied using brush strokes.*

#### Supplementary Note 2: SproutSAM results

##### Prompt Sampling Sensitivity Across Models

To evaluate the effect of prompt sampling strategies on segmentation accuracy, we tested seven configurations across five foundation models and three datasets (dog, foraminifera, and EM). The seven sampling configurations varied along three axes: sampling strategy (random, k-means, centre–edge), number of positive/negative points (e.g., 1 vs. 3), and whether negatives were sampled per class or from all other classes combined (shared negatives).

As shown in *Supplementary Fig. 6*, performance varied depending on the sampling strategy, particularly for ViT-L-em. In the EM dataset, ViT-L-em showed a large drop in performance when using the simplest random: 1 pos, 1 neg/class configuration, with Dice scores falling below 0.4. In contrast, k-means clustering and centre–edge strategies improved segmentation accuracy.

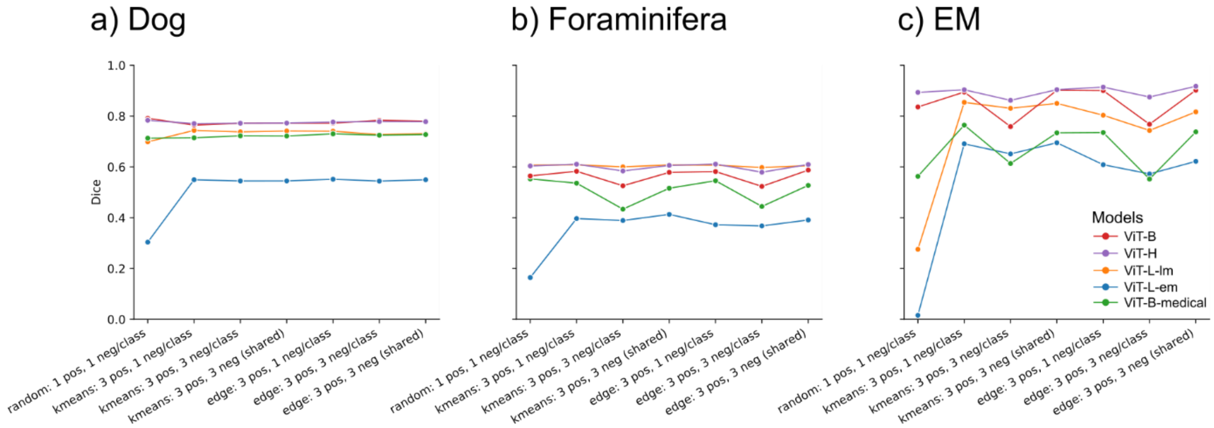


***Supplementary Figure 6. Effect of prompt sampling strategies on segmentation accuracy across datasets and models.*** *Each panel shows the Dice score of five segmentation foundation models (ViT-B, ViT-H, ViT-L-lm, ViT-L-em and ViT-B-medical) across seven different prompt sampling configurations, evaluated on three datasets: (a) Dog CT scans, (b) Foraminifera CT scans, and (c) Electron Microscopy of the mouse retina*.

To quantify the overall effects of sampling design, we computed the mean Dice score across all datasets and models for each configuration. The results revealed that k-means: 3 pos, 1 neg/class achieved the highest average performance (mean Dice = 0.656), followed closely by k-means: 3 pos, 3 neg (shared) and centre–edge: 3 pos, 3 neg (shared). In contrast, the random sampling strategy consistently underperformed (mean Dice = 0.543).

Together, these findings demonstrate that prompt sampling strategies have an influence on segmentation performance in SproutSAM, and that structured (non-random) strategies generally yield more stable and accurate results.

#### Examples of SproutSAM

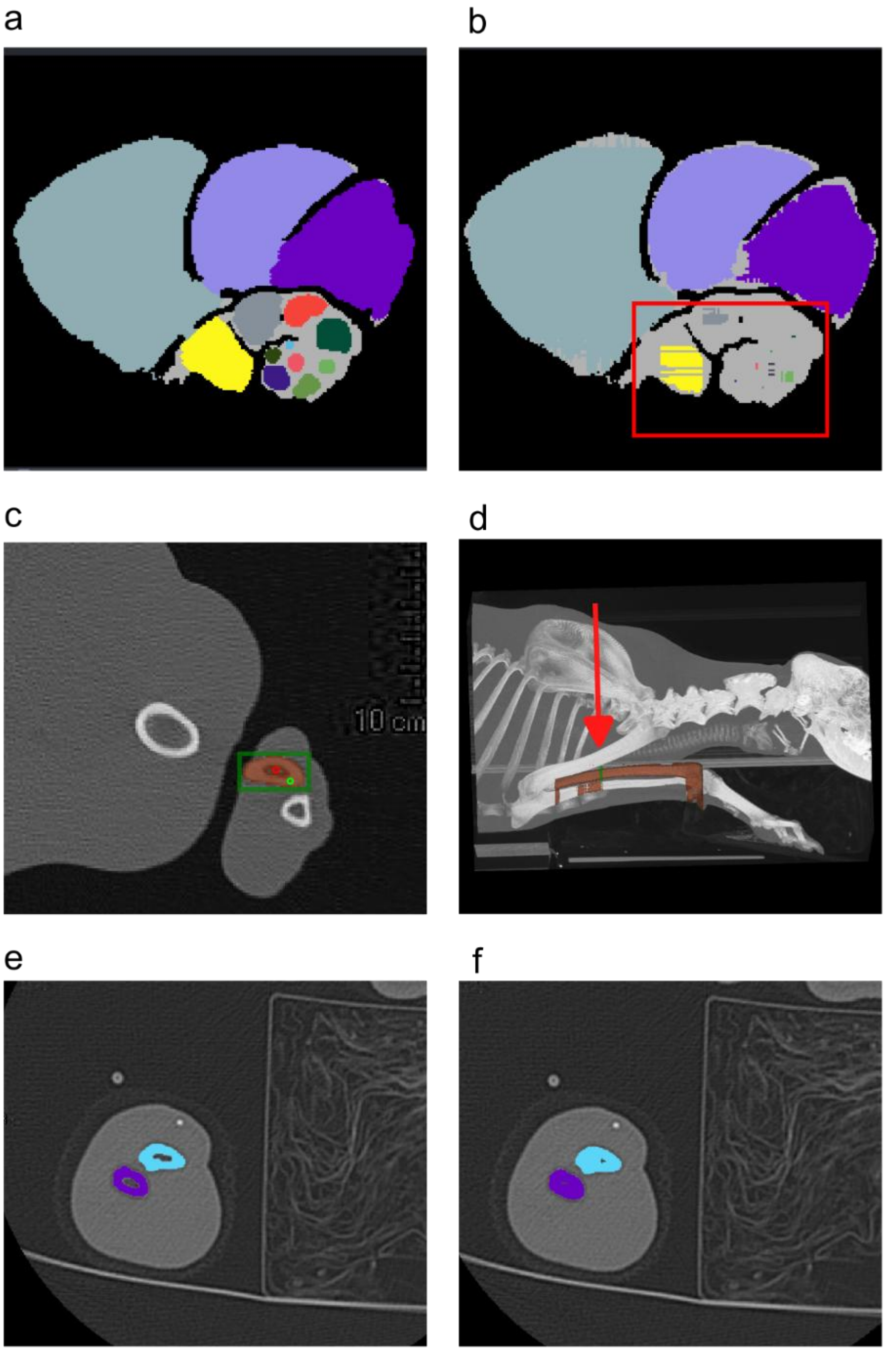


***Supplementary Figure 7. Representative segmentation failure cases in SproutSAM and MicroSAM.*** *(a) Ground truth segmentation of a CT slice containing multiple bone elements. (b) SproutSAM result on the same slice, showing under-segmentation and misclassification between adjacent structures (errors highlighted in red; zoomed-in view shown). (c) Example of MicroSAM prompts (bounding box and points) applied to a single slice. (d) 3D result propagated from (c), showing incomplete segmentations (cutoffs at both ends) and inclusion*

*of neighbouring structures. The slice with prompts is indicated by a red arrow. (e) Ground truth of a bone segmentation where internal cavities (e.g., medullary spaces) are correctly excluded. (f) An example of incorrectly segmented internal cavities, which can happen to both SproutSAM and MicroSAM. Zoom-in views in (a), (b) (c), (e), and (f) for better visualisation effect.*

#### Runtime

To further assess computational efficiency and scalability, we compared the runtime of SPROUT's Grow algorithm and SproutSAM using a set of 2D synthetic images and seeds (Supplementary Table 1). In each test case, a single-class seed was placed at the centre of a blank image. The initial seed was a 50-pixel square, and the target area was set to a square of 500, 1000, or 2000 pixels per side, depending on the test case.

Grow was applied with fixed dilation steps until full expansion (i.e., no further growth under threshold constraints) on a single CPU thread (no parallelisation). SproutSAM used points sampled from the seed and performed a single forward pass with the ViT-H model at 1024×1024 resolution (the default input size for SAM). While SproutSAM's runtime remained nearly constant (~8–9 s), Grow's runtime increased substantially with both image resolution and target mask size, reflecting its iterative, pixel-wise operations over the full image domain. In contrast, SproutSAM's runtime is determined primarily by the fixed-cost model inference, independent of mask size or image content, as long as resolution is standardised.

***Supplementary Table 1.*** *Runtime comparison of SPROUT's Grow algorithm and SproutSAM on 2D synthetic images. Each case starts from a 50-pixel seed at the centre of a blank image, expanding to the specified target area. Grow runtime increases with both resolution and target size, whereas SproutSAM runtime remains nearly constant.*

| Image Size | Grow Target Area | Grow Time | Dilation steps | SproutSAM Time |
|---|---|---|---|---|
| 512×512 | 500x500 pixels | ~3 s | 500 | ~8 s |
| 1024×1024 | 500x500 pixels | ~6 s | 500 | ~8 s |
| 1024×1024 | 1000x1000 pixels | ~30 s | 1000 | ~8 s |
| 2048×2048 | 500x500 pixels | ~35 s | 500 | ~9 s |
| 2048×2048 | 2000x2000 pixels | ~140 s | 2000 | ~9 s |

#### Supplementary Note 3: Evaluation of Parallelisation Efficiency

We assessed the computational performance of SPROUT's core functions: (i) seed generation, (ii) adaptive seed generation, and (iii) growth, with a focus on how multi-threaded parallelisation affects processing time for individual scans. The evaluation was conducted on three representative datasets: a CT scan of a French bulldog hindlimb, a binary segmentation of a *planktonic foraminifera*, and a micro-CT scan of an aardvark skull. Detailed scan characteristics and task configurations are summarised in Supplementary Table 2. All benchmarking was performed on a Windows 11 workstation equipped with an Intel Xeon Gold 5118 CPU @ 2.30 GHz (2 processors, 48 threads total). For each scan and task, we measured runtime using 1 (no parallelisation), 2, 4, 8, and 16 threads.

As shown in Supplementary Fig. 8, parallelisation significantly reduced runtime for adaptive seed and growth tasks across all scans. In the case of adaptive seed generation, increasing the thread count from 1 to 16 led to a minimum time reduction of 34%. For the growth step, the improvement was even more pronounced, with a minimum speedup of 68%, and up to 78% for scans with a large number of segmented regions, such as the French bulldog hindlimb and the aardvark skull.

In contrast, performance gains for seed generation were more limited. This is because parallelisation in this step only applies across threshold values, while most of the computational cost lies in morphological operations such as erosion and connected components analysis. Accordingly, speed improvements were only observed when the number of threads exceeded the number of threshold values (e.g., more than 11 threads for the French bulldog hindlimb scan, or more than 1 thread for the other two scans).

***Supplementary Table 2.*** *Scan details and task configurations are used for evaluating the parallelisation efficiency.*

| Object | Specimen ID | Bits | Dimension (XYZ) | Task Configurations |
|---|---|---|---|---|
| **French Bulldog left hindlimb** | 256334 | 8 | 182x241x1036 | **Seeds**: 11 thresholds, zero erosions and 30 segments. **Adaptive seed**: The same inputs as those used for seed generation. **Growth**: Five dilations on a list of 11 thresholds. |
| **Planktonic Foraminifera (*Menardella limbata*)** | st021_bl1_fo4 | 8 | 992x1015x180 | **Seeds**: One threshold, ten erosions and 25 segments. **Adaptive seed**: The same inputs as those used for seed generation. |

| | | | | |
|---|---|---|---|---|
| | | | | **Growth**: Ten dilations on one threshold. |
| **Aardvark skull (*Orycteropus afer*)** | AMNH 51909 | 8 | 1024x738x1237 | **Seeds**: One threshold, three erosions and 40 segments. **Adaptive seed**: The same inputs as those used for seed generation. **Growth**: Ten dilations on a list of four thresholds. |

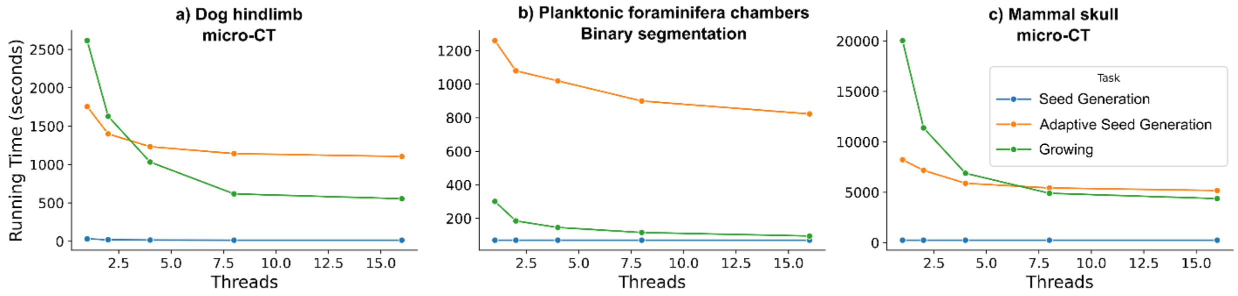


***Supplementary Figure 8.*** *Effect of thread count on running time for SPROUT functions across three representative datasets: (a) Dog hindlimb (CT), (b) Binary segmentation of planktonic foraminifera chambers, and (c) Mammal skull (micro-CT). Running time (in seconds) is plotted against the number of threads for three SPROUT functions: seed generation, adaptive seed generation, and growing.*

#### Scan detail

***Supplementary Table 3.*** *Summary of datasets used in this study, including object type, scan modality, source, intended usage, specimen identifiers, and scan dimensions*

| **Object** | **Scan type** | **Source** | **Usage** | **Specimen number** | **Dimension (XYZ)** |
|---|---|---|---|---|---|
| Dog | CT | University of Liverpool Small Animal Teaching Hospital | illustrative example | 228257(Labrador thorax) | 512x512x571 |
| | | | | 217408 (Dachshund right lower forelimb) | 448x406x226 |
| | | | | 255119 (Border Collie upper forelimbs) | 512x512x667 |
| | | | | 259296 (Dachshund hindlimbs and tail) | 256x256x961 |
| | | | • Interpolation | 200000 (Greyhound, left) | 257x371x580 |

| | | | | | |
|---|---|---|---|---|---|
| | | | • SproutSAM | 200000 (Greyhound, right) | 229x363x580 |
| | | | | 242343 (Border Collie, left) | 512x512x1182 |
| | | | | 242343 (Border Collie, right) | 512x512x1431 |
| | | | | 255544 (French Bulldog, right) | 512x512x581 |
| | | | | 256334 (French Bulldog, left) | 182x241x1036 |
| | | | | 256334 (French Bulldog, right) | 202x250x1036 |
| | | | | 257098 (Dachshund, left) | 210x512x672 |
| | | | | 257098 Right forelimb (Dachshund, right) | 199x512x638 |
| | | | Deep Learning training set | 241035 (French Bulldog) | 226x512x276 |
| | | | | 247854 (West Highland White Terrier) | 301x512x210 |
| | | | | 248812 (West Highland White Terrier) | 226x512x219 |
| | | | | 250482 (West Highland White Terrier) | 251x512x174 |
| | | | | 250882 (Border Collie) | 276x512x521 |
| | | | | 251908 (Great Dane) | 161x326x476 |
| | | | | 254861 (Beagle) | 126x256x319 |
| | | | | 257593 (Dachshund) | 136x226x151 |
| | | | | 258467 (Dachshund) | 111x256x201 |
| | | | | 259056 (West Highland White Terrier) | 176x512x649 |
| | | | Deep learning test set | 228420 (Dachshund) | 116x226x151 |
| | | | | 232558 (Beagle) | 126x256x299 |
| | | | | 238046 (Wolfhound) | 226x512x1151 |
| | | | | 239179 (Yorkshire Terrier) | 262x512x163 |
| | | | | 244195 (Labrador) | 236x512x749 |

| | | | | | |
|---|---|---|---|---|---|
| Planktonic foraminifera chambers | Binary mask | Ocean Drilling Program (ODP) Site 925, μ-VIS X-ray Imaging Centre, University of Southampton, UK | • SproutSAM<br>• illustrative example | st018_bl1_fo4 | 992x1015x108 |
| | | | | st021_bl1_fo4 | 992x1015x180 |
| | | | | st051_bl4_fo3 | 988x1011x163 |
| | | | SproutSAM | st050_bl1_fo3 | 988x1011x561 |
| | | | | st051_bl1_fo4 | 988x1011x232 |
| | | | | st061_bl1_fo2 | 988x1011x188 |
| | | | | st062_bl1_fo2 | 988x1011x133 |
| | | | | st049_bl1_fo2 | 992x1015x179 |
| | | | | st070_bl1_fo1 | 988x1011x234 |
| | | | | st098_bl1_fo5 | 980x1011x268 |
| Mouse retina | Electron microscopy | (Helmstaedter et al., 2013) | • Ilastik<br>• SproutSAM illustrative example | N/A | 256x256x256 |
| Human endothelial cells | 2D confocal microscopy | Laboratory for Vascular Morphogenesis, RIKEN Center for Biosystems Dynamics Research, Japan | illustrative example | N/A | 2796x2796 |
| *Papio hamadryas* skull | Micro-CT | American Museum of Natural History (downloaded from Morphosource) | illustrative example | AMNH:Mammals:M-52674 | 686x630x1179 |
| *Orycteropus afer* skull | Micro-CT | The University of Texas High-Resolution X-ray CT Facility & American Museum of Natural History (downloaded from Digimorph) | • illustrative example<br>Time Evaluation | AMNH 51909 | 1024x738x1237 |
| *Tiliqua scincoides* | Micro-CT | California Academy of Sciences & oVert TCN | illustrative example | CAS:HERP:254658 | 567x1626x2000 |
| Papuan weevil | Synchrotron CT | (Lösel et al., 2020) | illustrative example | N/A | 1497x734x1117 |
| Human heart | MRI | (Pace et al., 2015) | illustrative example | pat0 | 127x207x141 |
| Human Abdomen | CT | (Ma et al., 2022) | illustrative example | Tr_0001 | 110x512x512 |

| | | | | | |
|---|---|---|---|---|---|
| Steel reinforced concrete | Micro-CT | (Hampe, 2024) | illustrative example | N/A | 949x957x924 |

### Supplementary Method

#### Supplementary Note 4: SproutSAM Methods

SproutSAM adapts 2D segmentation foundation models (e.g., SAM) for volumetric segmentation by generating slice-wise prompts and predictions along the three axes (X, Y, and Z). Given an input segmentation containing one or more semantic classes, prompts are extracted from 2D slices sampled along each axis. For 2D inputs, no slicing is required; prompts are sampled directly from the 2D image, and the same fusion strategy applies. An overview of the SproutSAM workflow is illustrated in Supplementary Fig. 9.

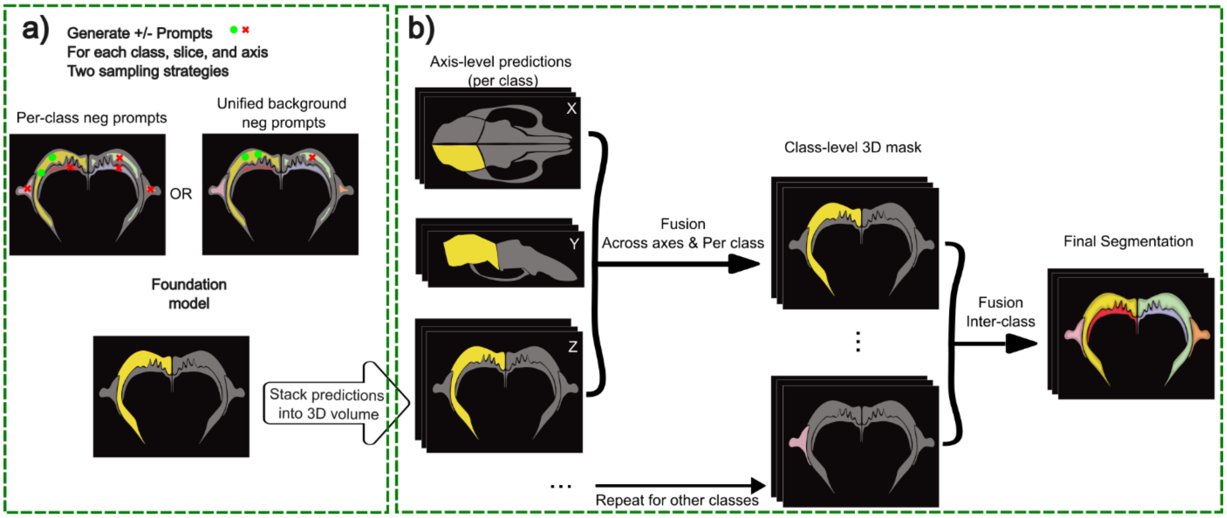


***Supplementary Figure 9.*** *(a) Prompt sampling strategies. SproutSAM generates positive (green) and negative (red) point prompts for each class, slice, and axis. Two strategies are illustrated: sampling negatives from each non-target class, or collectively from all non-background regions. Prompts are passed to a foundation model to produce axis-aligned segmentations. (b) Multi-axis and multi-class fusion. For each class, predictions along the X, Y, and Z axes are stacked into 3D volumes and fused via voxel-wise majority voting to produce a class-level 3D mask. This procedure is repeated for all classes. Final segmentation is obtained by resolving inter-class overlaps using a default or user-defined priority rule.*

##### Prompt Sampling Strategies

On each 2D slice, positive prompts are sampled from within the target region (i.e., the current semantic class), while negative prompts are sampled from regions belonging to other classes. Prompts are always sampled as 2D points, regardless of the model type (SAM or SAM2). Sampling methods include the following configurable options:

- Number of points sampled for positive and negative point prompts.
- Negative point prompts are sampled in one of two ways: either randomly from all other non-background regions, or separately from each non-target class.
- Point sampling strategies.
    - Random: Points are sampled uniformly from the foreground region.
    - K-means: Points are spatially distributed using k-means clustering to maximise coverage, especially useful for fragmented or irregular masks.
    - Centre-edge: One point is sampled from the centre of the region (maximum distance from the boundary), and the remaining points from near the edges to capture both core and boundary structure.

Prompt sampling is performed independently for every class and every slice. The resulting prompts are passed to the foundation model, which returns a binary mask prediction for each class–slice pair (Supplementary Fig. 9a illustrates the sampling strategies and example prompts used).

##### Multi-Axis and Multi-Class Fusion

After obtaining predicted 2D masks for each semantic class along all slices and axes, SproutSAM reconstructs full 3D segmentations using a two-stage fusion approach (Supplementary Fig. 9b):

1. Axis-wise intra-class fusion: For each class, masks predicted along the X-, Y-, and Z-axes are first stacked into three separate 3D volumes. These axis-aligned volumes are then combined using majority voting at the voxel level. A voxel is assigned to the class if at least two out of three axis volumes mark it as foreground.
2. Inter-class fusion: Once a 3D binary mask is generated for each class, the masks are merged into a single multi-class segmentation. Voxel conflicts (i.e., multiple classes claiming the same voxel) are resolved based on default class priority (ascending class index), or user-defined priority scheme.

#### Supplementary Note 5: napari-sprout interface and usage

To facilitate interactive parameter tuning and region-level editing, we developed napari-sprout, a plugin for the napari image viewer. The plugin enables users to load and visualise volumetric images and segmentations, run SPROUT functions via GUI, and perform semi-automated segmentation and post-processing.

Key features include:

- Interactive seed generation and growing with adjustable parameters (Supplementary Fig. 1a).
- Threshold preview, with per-class mask visualisation (Supplementary Fig. 1b).
- Region-level editing tools: merge, split, assign class labels (Supplementary Fig. 1c and d).
- Batch processing assistant: create CSV files for batch processing based on input folders.

To use napari-sprout, install napari and the plugin as follows:

```
pip install napari
cd ./napari_sprout
pip install -e .
```

You can launch napari by running *napari* in the command line. The napari-sprout plugin will appear under the Plugins menu in the napari UI. To activate the plugin, select the napari-sprout module from the plugin list.

The plugin is cross-platform and open-source, and regularly updated at the SPROUT repository: https://github.com/EchanHe/SPROUT. For the latest feature list and installation instructions, please refer to the README of the repository. While napari-sprout enables full GUI-based workflows for individual volumes, batch processing is currently supported via the script-based pipeline only (See Supplementary Note 8: Batch Processing Workflow)

#### Note 6: Other integrations Avizo and ImageJ

In addition to napari-sprout, we provide integrations with the widely used imaging platforms Avizo (Thermo Fisher Scientific, USA) and ImageJ. These tools are intended to support users who prefer working within familiar graphical environments when applying or refining results from the SPROUT workflow.

For Avizo, we developed a set of custom add-ons with graphical user interfaces (GUIs) that streamline the SPROUT workflow and reduce manual processing steps. These add-ons support key functions including pre-processing, batch import of SPROUT outputs, and manipulation of segmentation results (e.g., merging and splitting regions).

To extend compatibility to commonly used 2D datasets such as microscopy images, we also developed two ImageJ macros. These enable users to: (i) batch visualise SPROUT outputs; and (ii) manually edit segmentation masks using ImageJ's native annotation tools (e.g., the multi-point tool). This approach offers an intuitive and accessible interface for refining both seed masks and segmentation results, particularly for users working with 2D data in lightweight environments.

#### Note 7: Parallelisation and Memory optimisation

SPROUT supports parallel processing using Python's threading module. To enable this, users can specify the num_threads field in the YAML configuration. This significantly accelerates processing, especially for large 3D volumes or multiple class labels.

SPROUT utilises Python's threading module to support parallel processing at multiple levels. For batch processing, it enables the simultaneous processing of multiple images (file-level parallelisation). Furthermore, for each individual input, parallelisation is implemented for the seed generation, adaptive seed, and growth steps.

##### Memory Usage in Parallelisation

At the per-input level, memory usage has been optimised primarily by utilising sub-volumes for each segmentation class and removing unnecessary memory allocation during parallelisation. Here, we also describe the per-input memory usage for the primary SPROUT functions.

Without parallelisation, the memory usage is approximately $M = C \times S_{input}$. where $S_{input}$ represents the size of the input volume and $C$ is a constant. $C$ is 2 for *make_seeds.py* and *make_seeds_merged.py*, as these functions create a copy of the input volume for connected components or thresholding. For *make_grow_result.py*, $C$ is 3 due to the additional volume allocated for the candidate seed during the growth step.

With parallelisation, memory allocation varies depending on the function. For seed generation, which is parallelised based on thresholds, memory usage scales with the number of threads and is $M = C \times S_{input} \times T$, where $T$ is the number of threads. For adaptive seed generation and growth, which are parallelised based on classes, memory usage depends on the sub-volume size of each class. Memory usage is calculated as:

$$M = MAX(C \times S_{input} \times T, C \times S_{input} + \sum_{i=1}^{T} S_{classes,i})$$

Here $S_{classes} = \{S_{class1}, S_{class2}, \dots, S_{classm}\}$ is the list of sub-volume sizes of m classes.

#### Note 8: SPROUT algorithm modules and key parameters

This section provides an overview of how to run SPROUT using its input configuration (.yaml) files, followed by a description of the key parameters for each algorithm module. The .yaml files define the inputs, processing parameters, and output settings in a reproducible and shareable format. For the most up-to-date usage instructions, detailed parameter descriptions, and example templates, please refer to the SPROUT GitHub repository (https://github.com/EchanHe).

##### Seed Generation

To generate initial seeds from a grayscale image, users can run the following command

```
python sprout.py --seeds --config path/to/seed_config.yaml
```

Key parameters include:

- img_path: path to a grayscale .tif/.tiff image
- thresholds: the lower threshold. Can be given as a single number or as multiple values.
- upper_thresholds: the upper threshold. it should be larger than the corresponding lower threshold, defining a range-based thresholding.
- erosion_steps: the number of erosion iterations to apply (zero or more)
- segments: keep the N largest components

Other configuration options, such as output folder location, number of threads, or file-naming settings, are described in the repository under template/ and docs/config_seed.md.

##### Adaptive Seed Generation

For more robust segmentation across structurally similar datasets, SPROUT supports adaptive seed generation, which explores a range of parameters.

This method progressively generates smaller regions at each step and compares them to the previous step's results. To ensure that each output is smaller than the one before, the algorithm proceeds either by incrementally narrowing the threshold range (threshold-based adaptive seeds) or by increasing the number of erosion iterations (erosion-based adaptive seeds). If a single threshold is provided, then it runs as an erosion-based adaptive seed generation, and if a list of thresholds is provided, it runs as a threshold-based adaptive seed generation

```
python sprout.py --adaptive_seed --config path/to/adaptive_seed_config.yaml
```

Key parameters include:

- img_path: path to a grayscale .tif/.tiff image
- thresholds: the lower threshold. Can be given as a single number or as multiple values.
- upper_thresholds: the upper threshold. it should be larger than the corresponding lower threshold, defining a range-based thresholding.
- erosion_steps: the number of erosion iterations to apply (zero or more)
- segments: keep the N largest components

Other configuration options—such as the output folder location, number of threads, boundary mask path, file-naming settings, split criteria, and rules for retaining small components—are described in the repository under docs/config_adaptive_seed.md.

When using threshold-based adaptive seeds, each successive threshold range should be set to become progressively narrower (i.e. the lower threshold increases and/or the upper threshold decreases with each step) to ensure that the resulting regions become smaller at each iteration.

#### Growth

To recover full structures from seed masks, the region growth module expands each region based on intensity thresholds.

```
python sprout.py --grow --config path/to/grow_config.yaml
```

Key parameters include:

- img_path: path to a grayscale .tif/.tiff image
- seg_path: path to the seed/segmentation image for growing
- thresholds: the lower threshold. Can be given as a single number or as multiple values.
- upper_thresholds: the upper threshold. it should be larger than the corresponding lower threshold, defining a range-based thresholding.
- dilation_steps: the number of dilation iterations to apply (zero or more)

Other configuration options—such as the output folder location, number of threads, boundary mask path, file-naming settings and specific regions to grow—are described in the repository under template/ and docs/config_grow.md.

When using multiple threshold ranges for the growth step, each successive range should be set wider than the previous one (i.e. lowering the lower threshold and/or raising the upper threshold) to allow the grown regions to progressively recover more of the target structure.

#### SproutSAM

SPROUT also supports the use of segmentation foundation models (e.g., SAM, SAM2) to convert seed masks into final segmentations.

```
python sprout.py --sam --config path/to/SproutSAM_config.yaml
```

Key parameters include:

- img_path: path to a grayscale .tif/.tiff image
- seg_path: path to the seed/segmentation image for growing
- n_points_per_class: Number of positive points to sample from each class.
- negative_points: Number of negative points to sample for each class.
- sample_method: Method to sample point prompts

Other configuration options, such as the output folder location, are described in the repository under template/ and docs/config_sam.md.

#### Batch Processing Workflow

SPROUT supports batch processing for all core modules, including seed generation, adaptive seed generation, growing, and SproutSAM.

Batch mode can be enabled using the *--batch* flag in CLI. For example:

```
python sprout.py --seeds --batch --config path/to/batch_seed.yaml
```

Each batch run requires a CSV file specifying the input parameters (e.g., paths to input images or seed masks) and a YAML configuration file that defines global settings (e.g., threshold values, erosion steps, and number of threads). Additionally, per-input customisation is supported via the CSV file. For instance, users may apply adaptive thresholds such as Otsu values computed from individual inputs or adjust erosion steps based on image resolution. This flexibility enables fine-tuned control for heterogeneous datasets, including specimens with varying sizes or imaging artefacts. All batch operations automatically generate log files that record segmentation statistics, runtime performance, and any encountered errors, allowing users to monitor progress and troubleshoot failures efficiently.

To assist users in preparing batch CSVs, napari-sprout includes a CSV Maker interface, which provides an interactive way to generate input tables based on file directories and user-specified options.

#### Note 9: Software Requirements and Installation Guide

SPROUT is implemented in Python (v3.10) and relies on a set of well-supported scientific computing libraries, including:

- NumPy (v1.26.4)
- Pandas (v2.2.1)
- Scikit-image (v0.22.0)
- Tifffile (v2024.2.12)
- PyYAML (v6.0.1)
- Trimesh (v4.3.1)
- Open3D (v0.18.0)
- Matplotlib (v3.8.3)

The graphical user interface napari-sprout, additionally requires:

- napari (v0.4.19 or later)
- magicgui and napari-tools-menu for GUI integration

The SAM-based segmentation module requires:

- PyTorch (v2.1 or later)
- segment-anything
- SAM2

SPROUT has been tested on Windows 10/11, Ubuntu 24.04 LTS, and MacOS 15, and supports multi-threaded execution, making it suitable for high-performance computing (HPC) clusters and cloud platforms such as AWS EC2. Plugins for Avizo (tested with Avizo 2020, 2022 Pro, 2023 Pro) and ImageJ (tested with ImageJ 1.54f) are also included.
